# Tyrosine kinase inhibitors affect sweet taste and dysregulate fate selection of specific taste bud cell subtypes via KIT inhibition

**DOI:** 10.1101/2025.08.15.670608

**Authors:** Christina M. Piarowski, Jennifer K. Scott, Courtney E. Wilson, Heber I. Lara, Ernesto Salcedo, Andrew S. Han, Elaine T. Lam, Peter J. Dempsey, Jakob von Moltke, Linda A. Barlow

## Abstract

Taste dysfunction, or dysgeusia, is a common side effect of many cancer drugs. Dysgeusia is often reported by patients treated with anti-angiogenic tyrosine kinase inhibitors (TKIs), which inhibit receptor tyrosine kinases (RTKs). However, the mechanisms by which TKIs cause dysgeusia are not understood, as the role of RTKs in adult taste homeostasis is unknown. Here, we find that treating adult mice with the TKI cabozantinib shifts the fate of differentiating functional taste cell subtypes within taste buds. Through behavioral assays, we find this cell fate shift leads to blunted responses to sweet in cabozantinib-treated mice. Finally, we show that inducible knockout of the RTK KIT, which is inhibited by cabozantinib, phenocopies taste cell fate shifts induced by TKI treatment. Our results establish KIT as a regulator of taste cell homeostasis and suggest that KIT inhibition may underlie TKI-induced dysgeusia in patients.

**Summary:** KIT signaling blockade by tyrosine kinase inhibitors alters the fate of functional taste bud cell subtypes.

## Introduction

The sense of taste, or gustation, plays a central role in how animals experience the world. Taste conveys important nutritional information through five modalities: sweet, bitter, umami, sour, and salty (Liman et al., 2014). Taste dysfunction (dysgeusia) is a common side effect of many cancer therapies (Gaillard and Barlow, 2021), which leads to significantly reduced quality of life, including weight loss and depression, as well as worsened long-term outcomes for patients (Epstein and Barasch, 2010, Sánchez-Lara et al., 2010). Improving both quality of life and treatment outcomes therefore necessitates interventions that mitigate or prevent dysgeusia. However, the field lacks important mechanistic understanding of how most cancer therapies affect the taste system (Gaillard and Barlow, 2021).

One class of targeted cancer therapies that commonly cause dysgeusia are anti-angiogenic tyrosine kinase inhibitors (TKIs). These TKIs reduce tumor vascularization by inhibiting vascular endothelial growth factor receptors (VEGFRs) and platelet-derived growth factor receptor beta (PDGFRβ), both receptor tyrosine kinases (RTKs) (Vigarios et al., 2017, Kiselyov et al., 2007, Raica and Cimpean, 2010). A subset of anti-angiogenic TKIs, including cabozantinib, axitinib and sunitinib, are approved as first- and second-line therapies for metastatic renal cell carcinoma (mRCC) and cause dysgeusia in 10-50% of mRCC patients (Vigarios et al., 2017, Hahn et al., 2019, Roberto et al., 2021, Hamazaki and Uesawa, 2024). However, the intended targets, VEGFRs and PDGFRβ, are rarely found in epithelial cells, as they are expressed primarily in endothelial and stromal cells (Strell et al., 2024, Koch et al., 2011). Further, transcriptome profiling has not detected VEGFR or PDGFRβ mRNAs in murine taste epithelium (Schaum et al., 2018, Sukumaran et al., 2017, Ren et al., 2017, Yamada et al., 2021, Golden et al., 2021, Shechtman et al., 2023, Lee et al., 2017). Cabozantinib, axitinib, and sunitinib, as well as TKIs more generally, inhibit many off-target RTKs including RET, MET, KIT, and others (Klaeger et al., 2017, Karaman et al., 2008, Davis et al., 2011). Many of these off-target RTKs are expressed in taste epithelium (Biggs et al., 2016, Choo and Dando, 2021, Donnelly et al., 2018, Mclaughlin, 2000, Sukumaran et al., 2017, Yamada et al., 2021, Wang et al., 2025), raising the possibility that inhibition of one or more off-target RTKs underlies TKI-induced dysgeusia.

Taste is mediated by taste buds, which are collections of 50-100 specialized epithelial taste bud cells (TBCs) categorized into 3 morphological types: type I, type II and type III (Delay et al., 1986, Kinnamon et al., 1993). Type I cells are glial-like support cells, type II cells detect sweet, bitter, or umami tastants, while type III cells detect sour and some salty tastants (Liman et al., 2014, Finger and Silver, 2000, Ohtubo and Yoshii, 2011, Ogata and Ohtubo, 2020). Taste buds also house specialized sodium-sensitive cells similar to type II cells in morphology, developmental regulation, and expression of many type II cell markers (Chandrashekar et al., 2010, Nomura et al., 2020, Ohmoto et al., 2020). All TBCs are short-lived, with lifespans ranging from 10 to 40 days, and are steadily generated by proliferating progenitors outside of buds (Perea-Martinez et al., 2013, Beidler and Smallman, 1965, Farbman, 1969)(and see (Barlow, 2015, Piarowski et al., 2025) for review). Despite this continual turnover, the proportions of TBC types are believed to be relatively stable throughout life (Barlow, 2015).

Thus, the renewal process, spanning progenitor proliferation to TBC fate specification, differentiation, and survival, must be tightly regulated to ensure reliable taste function over time. As TKIs used to treat mRCC are frequently associated with taste dysfunction (Hamazaki and Uesawa, 2024, Vigarios et al., 2017), characterizing how these drugs impact TBC homeostasis may provide cellular and molecular mechanistic insight into how this class of TKIs causes dysgeusia.

Taste buds are housed in specialized gustatory papillae on the tongue. In rodents, fungiform papillae (FFP) are distributed throughout the anterior tongue and contain one taste bud each, while in the posterior tongue, bilateral foliate papillae and a single circumvallate papilla (CVP) at the posterior midline both comprise invaginated epithelial trenches that house hundreds of taste buds (Kinnamon, 1991). Importantly, taste buds in all papillae detect sweet, sour, bitter, etc., although bitter sensitivity is greater in the CVP and sweet sensitivity is greater in anterior FFP (Shingai and Beidler, 1985, Ninomiya et al., 1993). Additionally, specialized sodium-sensing type II-like cells are found only in FFP (Chandrashekar et al., 2010). How TKIs impact taste buds, and whether their impact differs between taste fields, is not known.

Here, we investigated the effects of anti-angiogenic TKIs on taste homeostasis using lingual organoids and mouse models. We show that in the CVP, cabozantinib affects renewal of type II TBC subtypes by dysregulating fate selection of differentiating type II cells such that production of sweet TBCs is diminished and that of bitter/umami cells is increased. Changes in type II TBC composition in CVP taste buds corresponds with blunted sweet taste behavior, thus linking dysregulation of type II TBC fate selection with dysgeusia. Because sweet sensitivity is greater in FFP, we were surprised to find that drug treatment did not block sweet TBC differentiation and instead reduced sodium-sensing type II cells in FFP. Finally, we find that the off-target RTK KIT likely regulates renewal of type II TBC subtypes in both CVP and FFP, as inducible *Kit* knockout in the type II TBC lineage phenocopies TKI treatment. This work provides insight into the regulation of type II TBC subtype fate selection, establishes KIT signaling in this process, and implicates KIT inhibition as a driver of TKI-induced dysgeusia.

## RESULTS

### TKIs selectively affect sweet type II TBCs in lingual organoids

We first used lingual organoids to screen the effect of TKIs on taste bud homeostasis, as organoids generated from GFP^+^ taste progenitors from the CVP epithelium of adult *Lgr5^CreER-eGFP^* mice contain all TBC types, non-taste epithelial cells and progenitors (Shechtman et al., 2021, Ren et al., 2014). We assessed if three TKIs used in treatment of mRCC, cabozantinib, axitinib, and sunitinib (Roberto et al., 2021, Hahn et al., 2019), hindered cell proliferation and/or organoid survival during the growth phase of culture (days 2-6) (**Figure S1A**). To quantify proliferation, 5-ethynyl-2’-deoxyuridine (EdU) was added to cultures for 30 minutes prior to harvest; we found that TKI treatment did not alter EdU incorporation by organoids (**Figure S1B-C**). We further assessed organoid growth by measuring organoid size at day 6. As a positive control, organoids were treated with paclitaxel, which blocks cell proliferation (Weaver, 2014). These organoids were significantly smaller than negative controls, whereas TKI treatment had little to no impact on organoid growth. Organoids treated with 100nM axitinib were significantly smaller, but this difference was small compared to the impact of paclitaxel (**Figure S2A**). Results with a Cell-Titer Glo® 3D assay, which measures ATP luminescence as a proxy for cell survival, were congruent with organoid size; paclitaxel decreased luminescence compared to controls, whereas TKIs, including 100nM axitinib, had no impact (**Figure S2B**). Lastly, we quantified organoid survival at day 6 and found 40-60% of isolated *Lgr5*-GFP^+^ progenitors formed organoids regardless of condition (**Figure S2C-D**). In sum, our data indicate that cabozantinib, axitinib, and sunitinib do not affect taste progenitor proliferation or survival *in vitro*.

Next, to test if TKIs affect TBC differentiation, we treated organoids during the differentiation phase of culture (days 6-12) (**Figure 1A**). Organoids were harvested for immunofluorescence (IF) for markers of type I (NTPDase2^+^) (Bartel et al., 2006), type II (PLCβ2^+^) (Clapp et al., 2004), and type III (CAR4^+^)(Chandrashekar et al., 2009) TBCs. The prevalence of TBC types did not differ in TKI-treated versus control organoids (**Figures 1B-C**, **S3A-D**). Type II TBCs comprise functional subsets, and those in the CVP that detect bitter or umami express the G-protein GUSTDUCIN (GUST) (Kim et al., 2003). However, GUST^+^ TBC prevalence was also unaffected by TKI treatment (**Figure 1D-E**). Thus, TKIs did not impact differentiation of type I, II, III, or bitter/umami type II TBCs *in vitro*.

**Figure 1:**
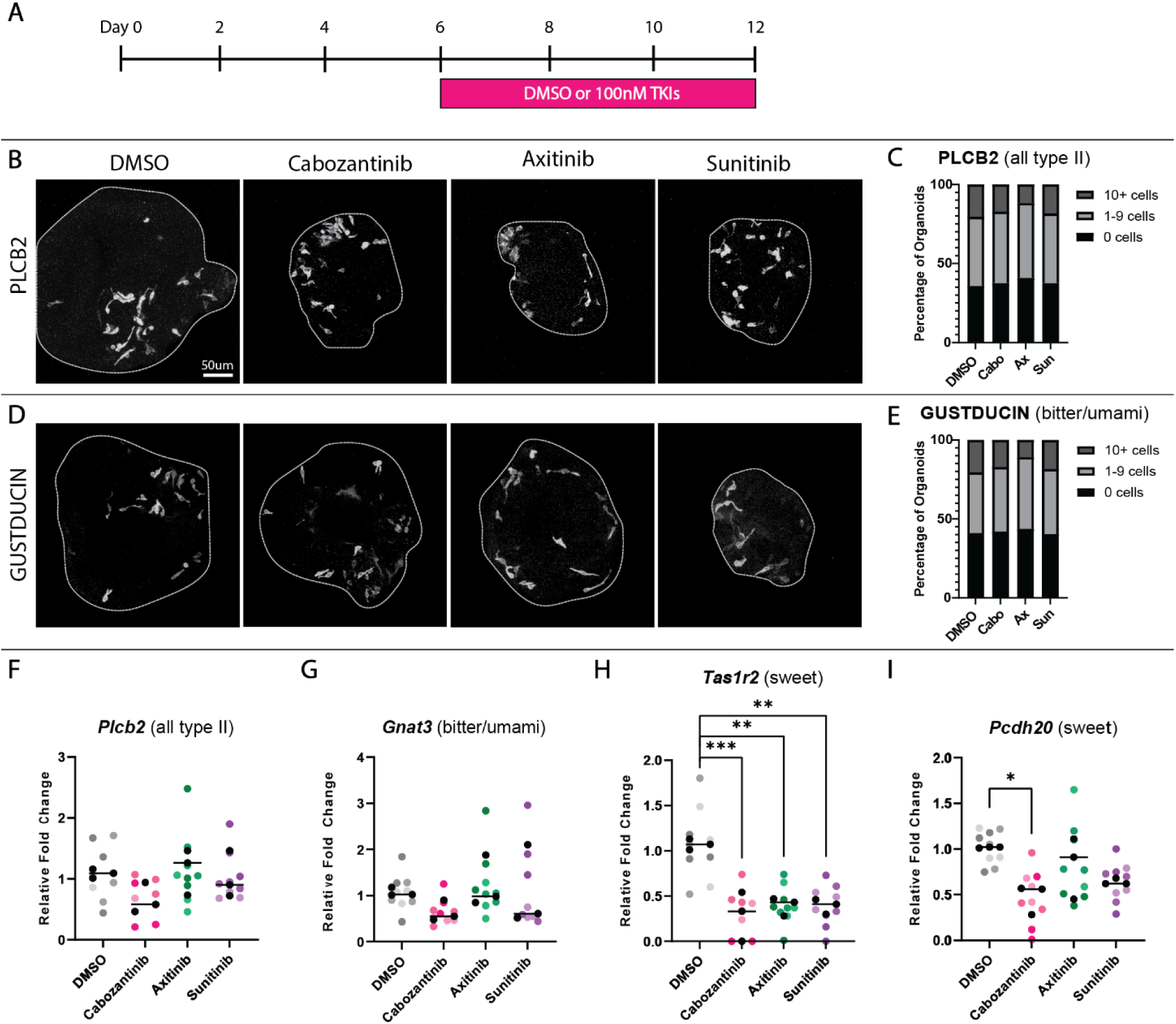
TKIs selectively reduce expression of sweet type II TBC markers in organoids. (**A**) Experimental design to test cabozantinib (Cabo), axitinib (Ax), and sunitinib (Sun) on TBC differentiation in organoids. (**B-E**) Compressed confocal z-stacks of control and TKI-treated organoids immunostained for markers of type II TBCs, PLCβ2 (**B**) and bitter/umami type II TBCs, GUST (**D**). Percentages of organoids containing 0, 1-9, or ≥ 10 cells immunopositive for PLCβ2 (**C**) or GUST (**E**) do not differ with treatment. Total organoids from 3 independent experiments: DMSO - 112; Cabo - 93; Ax - 110; Sun - 109. (**F-I**) Relative fold change in type II TBC marker expression via RT-qPCR. Each colored dot represents an individual sample of organoids pooled from 3 culture wells (over 200 organoids per sample). Within a condition, different shades represent 3 biological replicates and black dots are averages of each replicate. Ordinary one-way ANOVA with Dunnett’s multiple comparisons test (**F-G**) or Holm-Sidak’s multiple comparisons test (**I**) were used to compare average values (black line) for each marker across conditions (* p≤0.05, ** p≤0.01, *** p≤0.001).

To determine if TKIs affect other taste epithelial cell populations in organoids, we expanded our panel of markers using RT-qPCR. Expression of *Kcnq1*, expressed by all TBCs (Wang et al., 2009), and *Krt13*, a marker of non-taste CVP epithelium (Iwasaki et al., 2011), was unchanged by drug treatment, suggesting that production of taste versus non-taste lineages was unaffected by TKIs (**Figure S3E-F**). Consistent with immunostaining, markers of type I (*Kcnj1*) (Dvoryanchikov et al., 2009), type II (*Plcβ2*), and type III (*Pkd2l1*) (Kataoka et al., 2008) TBCs, as well as bitter/umami TBCs (*Gnat3* - encodes GUST) were unaffected by TKIs (**Figures 1F-G**, **S3G-H**). Surprisingly, we found that markers of the sweet subset of type II TBCs were specifically reduced. Expression of the G-protein coupled receptor *Tas1r2,* which heterodimerizes with TAS1R3 to mediate sweet transduction (Nelson et al., 2001), was decreased by all three TKIs (**Figure 1H).** *Pcdh20*, another sweet cell marker (Hirose et al., 2020), was also reduced significantly by cabozantinib and expression trended downward with sunitinib (p=0.07) (**Figure 1I).** These data suggest that TKIs selectively affect sweet type II TBCs, but not other cell types, in lingual organoids.

### Cabozantinib changes the cellular composition of taste buds *in vivo*

Since sweet TBC gene expression was reduced by TKIs *in vitro*, we asked if TKI treatment specifically affected sweet type II TBCs in the CVP of adult mice. Cabozantinib was selected because of its long half-life (∼120 hours) (Lacy et al., 2017) and because it significantly reduced expression of both sweet TBC markers tested in organoids (**Figure 1**). Mice were gavaged with vehicle or cabozantinib for 2 weeks (daily) or 4 weeks (5 days/week) (**Figure 2A-B**). Cabozantinib was well-tolerated by the mice, whose weights were stable over 4 weeks (**Figure S4A**). Taste buds were also grossly unaffected; taste bud number and size did not differ with treatment (**Figure S4B-I**). These data suggest that cabozantinib does not affect progenitor cell output or overall renewal of taste buds, consistent with data from organoids (**Figures S1-S2**).

**Figure 2:**
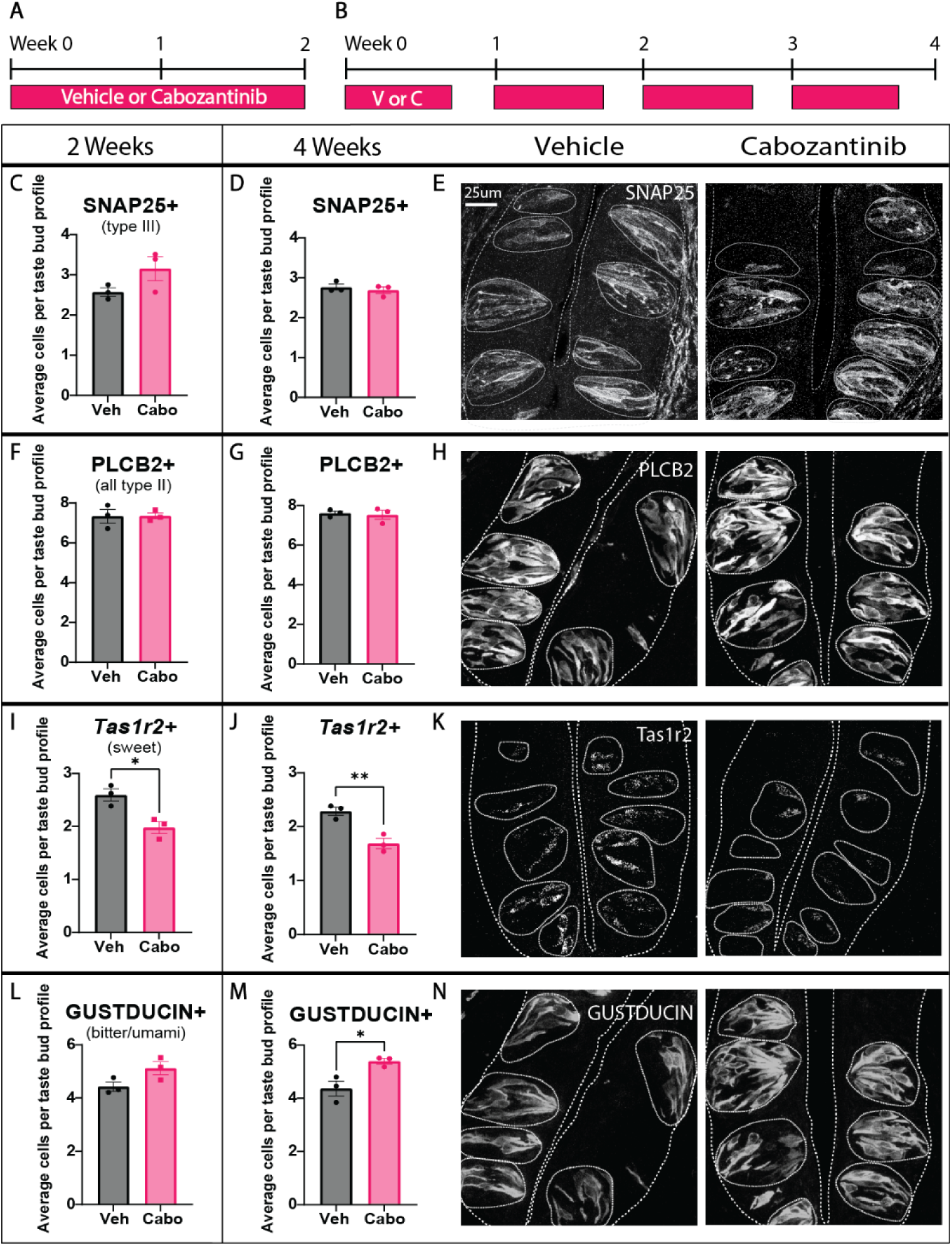
Cabozantinib changes the composition of type II TBC functional subtypes in CVP taste buds. Mice were treated daily for 2 weeks (**A**) or 5 days per week for 4 weeks with drug or vehicle (**B**). (**C-N**) Quantification of the average number of stained cells per taste bud profile with corresponding representative images for SNAP25 immunofluorescence (IF) (**C-E**), PLCβ2-IF (**F-H**), *Tas1r2* HCR *in situ* hybridization (**I-K**) and GUST-IF (**L-N**). Representative images are compressed confocal z-stacks. Coarse dashed lines delineate basement membrane and apical epithelial surface; fine dashed lines encircle individual taste buds. Scale bar in E applies to H, K, and N. In all histograms, each dot represents the average taste cell tally from each mouse (N=3 per condition, ∼80 taste buds/mouse). Unpaired t-test. Mean +/- SEM (* p≤0.05, ** p≤0.01).

To investigate cellular changes within taste buds, we first assessed the prevalence of type II (PLCβ2^+^) and type III (SNAP25^+^) (Yang et al., 2000) TBCs in drug-treated versus control mice. As expected from results in organoids, cabozantinib did not alter the number of type II or III cells (**Figure 2C-H).** Consistent with reduced *Tas1r2* expression in organoids, however, drug treatment significantly diminished the number of *Tas1r2^+^* sweet TBCs *in vivo* (**Figure 2I-K**). In contrast to organoids, however, where GUST^+^ type II TBCs were unaffected, cabozantinib increased GUST^+^ type II cells in CVP taste buds (**Figure 2L-N**). Thus, while the total number of type II cells was unaffected, TKI treatment shifted type II TBC subtype composition; *Tas1r2*^+^ sweet cells were reduced with a concomitant increase in GUST^+^ bitter/umami cells. Importantly, this phenotype was progressively robust with longer drug treatment (**Figure 2**), suggesting that the shift in type II subtypes depends on gradual turnover of TBCs.

*In vivo* TKI treatment revealed an increase in GUST^+^ bitter/umami cells not seen in organoids. Upon closer inspection, we found that GUST^+^ type II cells were over-represented in CVP-derived organoids compared to their proportion in CVP taste buds (80% vs 60%, respectively, of PLCβ2^+^ type II cells are GUST^+^) (**Figure S5A-B**). Thus, over-representation of GUST^+^ TBCs in organoids may have obscured any smaller increases in this cell population caused by drug treatment.

### Cabozantinib does not shift cell type composition via cell death or transdifferentiation

As cabozantinib significantly reduced *Tas1r2*^+^ sweet cells and increased GUST^+^ bitter/umami cells in the CVP, we reasoned this shift could be due to: (1) decreased sweet TBC survival, (2) transdifferentiation of sweet cells into bitter/umami cells, or (3) a shift in fate of newly differentiated type II cells. A previous study revealed that the RTK KIT is expressed by TAS1R3^+^ TBCs, which encompass sweet and umami type II TBC subtypes in CVP (Choo and Dando, 2021). More recently, KIT has been identified as a specific marker of sweet cells (Ki et al., 2025). Using hybridization chain reaction (HCR) *in situ* hybridization, we confirmed that *Kit* co-localizes with *Tas1r2* in CVP taste buds (**Figure S6A**). Consistent with co-expression of *Tas1r2* and *Kit*, and a decrease in *Tas1r2^+^* cells in drug-treated mice (see **Figure 2**), cabozantinib significantly decreased the number of KIT^+^ TBCs per CVP taste bud profile (**Figure S6B-D**). These results support previous findings that *Kit* is expressed by *Tas1r2^+^* sweet type II cells in CVP taste buds.

To assess the fate of differentiated sweet cells following drug treatment, we induced lineage tracing in *Kit^CreER/+^;Rosa26^YFP/YFP^* mice to label existing sweet cells and then treated animals with vehicle or drug for 2 weeks (**Figure 3A**). If cabozantinib decreases sweet cell survival, then we would expect fewer lineage traced *Kit*-YFP^+^ cells with drug treatment. However, *Kit-*YFP^+^ cell number did not differ in vehicle-versus cabozantinib-treated mice, indicating no impact on sweet cell survival (**Figure 3B-C**). We next examined type II subtype identity of these *Kit*-YFP^+^ cells to determine if cabozantinib caused transdifferentiation of differentiated sweet cells, potentially into GUST^+^ bitter/umami cells. Consistent with HCR results, in controls most *Kit*-YFP^+^ cells were PLCβ2^+^ type II cells (**Figure 3D-E**). Of these, ∼85% were *Tas1r2*^+^ sweet cells with very few GUST^+^ bitter/umami cells (**Figure 3F-I**), suggesting that *Kit* is expressed in a small subset of differentiated bitter/umami cells, and/or in cells that give rise to both sweet and bitter/umami cells. Regardless, cabozantinib did not change the percentage of *Kit*-YFP^+^ cells expressing *Tas1r2* or GUST (**Figure 3F-I**), demonstrating that differentiated sweet cells in the CVP maintain their identity and do not transdifferentiate into bitter/umami type II TBCs with drug treatment.

**Figure 3:**
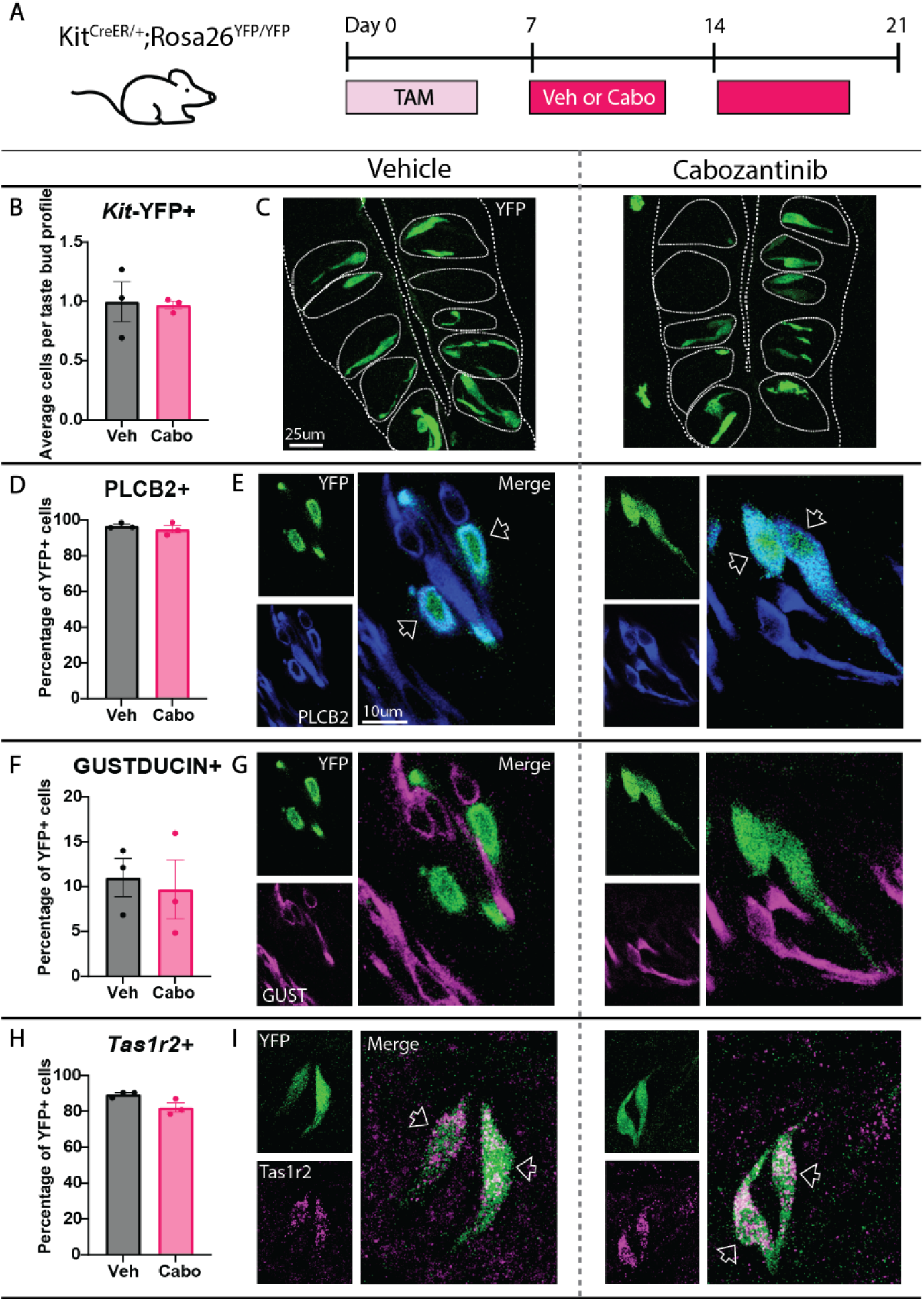
Cabozantinib does not impact survival or induce transdifferentiation of sweet type II TBCs in CVP taste buds. (**A**) *Kit^CreER/+^;Rosa26^YFP/YFP^* mice were gavaged with tamoxifen daily for 5 days. Following a 3-day chase with daily cage changes, mice were gavaged with vehicle or cabozantinib 5 days per week for 14 days. (**B**) *Kit*-YFP^+^ cells per taste bud profile were unaltered by TKI treatment. (**C**) Compressed confocal z-stacks of *Kit*-YFP^+^ taste cells in CVP taste buds from vehicle-versus cabozantinib-treated mice. Coarse dashed lines delineate basement membrane and apical epithelial surface, fine dashed lines encircle individual taste buds. (**D-I**) The percentage of *Kit*-YFP^+^ cells (green in all panels) co-expressing PLCβ2 (**D-E, blue in E,** optical section), GUST (**F-G, Magenta in G**, optical section), or *Tas1r2* (**H-I, magenta in I,** compressed z-stack), is unchanged by drug treatment. Empty arrowheads in **E** and **I** indicate double-labeled cells. In all panels, values were calculated across >200 YFP^+^ cells and >200 taste buds per condition. Unpaired t-test performed for all quantifications. Mean+/-SEM.

### Cabozantinib alters the fate of differentiating type II TBC subtypes

We next asked if TKI treatment alters the fate of differentiating TBCs. Sonic Hedgehog (SHH) marks post-mitotic taste precursor cells that differentiate into each of the TBC types. Precursor cells enter taste buds and express *Shh* transiently (for ∼24 hrs), then differentiate into TBCs within 3 days (Miura et al., 2014, Miura et al., 2006). We thus employed *Shh^CreER/+^;Rosa26^tdTomato/+^* mice to track the fate of new TBCs that differentiate during drug treatment (**Figure 4A**).

**Figure 4:**
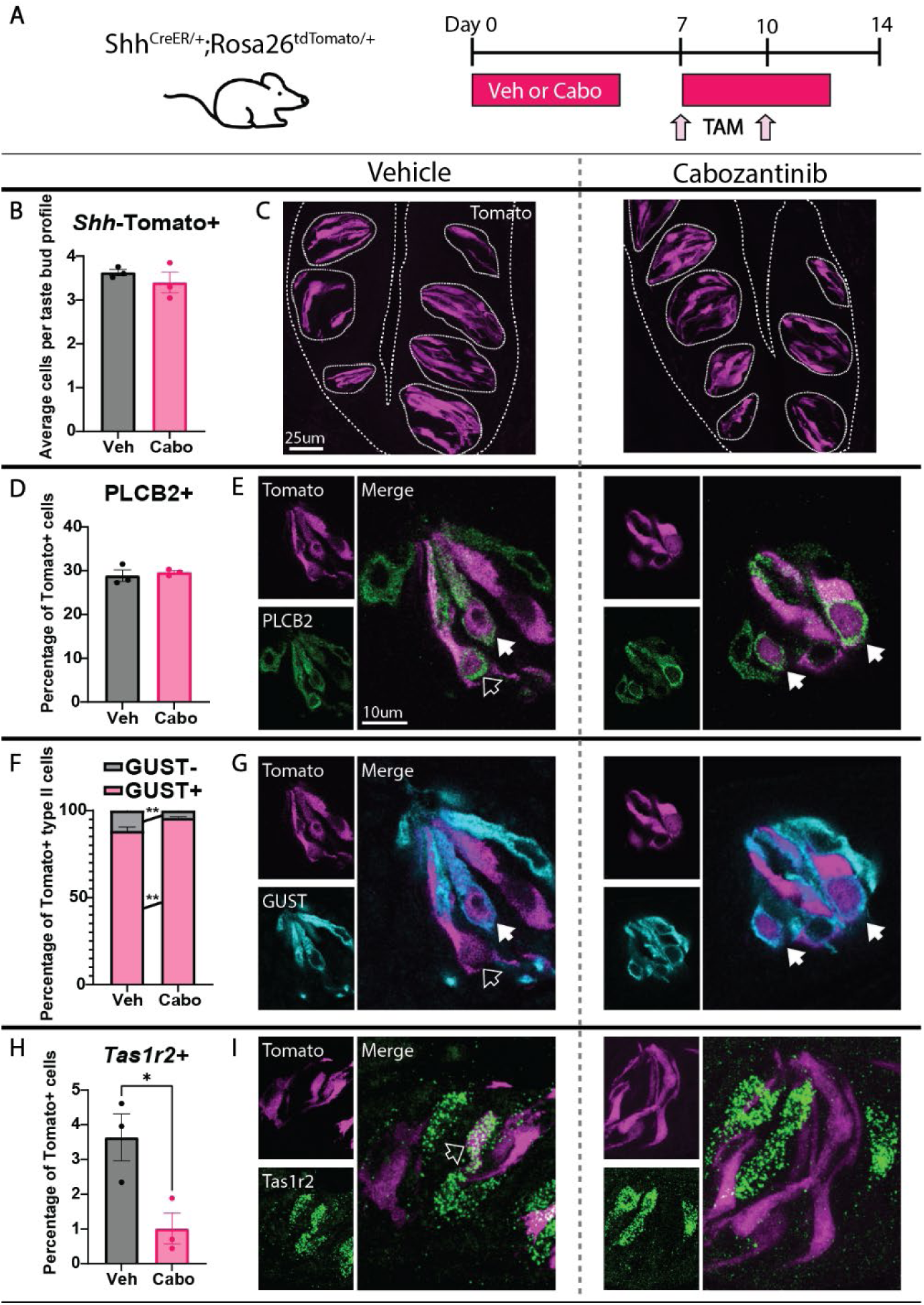
Cabozantinib shifts proportions of differentiating type II TBC subtypes in CVP taste buds. (**A**) *Shh^CreER/+^;Rosa26^tdTomato/+^* mice were treated with vehicle or cabozantinib 5 days per week for 14 days and dosed with tamoxifen on days 7 and 10 to label *Shh*^+^ taste precursor cells. (**B**) The number of *Shh*-Tomato^+^ cells per taste bud profile was unaltered by cabozantinib. (**C**) Compressed confocal z-stacks of *Shh*-Tomato^+^ TBCs in taste buds from vehicle-versus cabozantinib-treated CVPs. Coarse dashed lines delineate basement membrane and apical epithelial surface, fine dashed lines encircle individual taste buds. (**D-I**) The percentage of *Shh*-Tomato^+^ cells (**magenta** in all panels) co-expressing PLCβ2 did not change (**D-E, green in E,** optical section). The percentage of Tomato^+^/PLCβ2^+^ cells expressing GUST significantly increased, while the GUST^-^ percentage significantly decreased (**F-G, cyan in G,** optical section). The percentage of *Shh*-Tomato^+^ cells expressing *Tas1r2* was significantly decreased (**H-I, green in I,** compressed z-stacks). Empty arrowheads in **E**, **G** and **I** indicate double-labeled cells, white arrowheads in **E** and **G** indicate triple-labeled cells. In all panels, values were calculated across >400 Tomato^+^ cells and >200 taste buds per condition. Unpaired t-test (**B,D,H**) and two-way ANOVA with Sidak’s multiple comparisons test (**F**) were performed to compare average values. Mean +/- SEM (* p≤0.05, ** p≤0.01).

In the CVP, the number of *Shh*-Tomato^+^ TBCs was unaffected by TKI treatment, indicating cabozantinib does not impede overall differentiation of TBCs from *Shh*^+^ precursor cells (**Figure 4B-C**). Cabozantinib also did not affect differentiation of type II TBCs broadly, as ∼30% of *Shh*-Tomato^+^ cells were PLCβ2^+^ regardless of treatment (**Figures 4D-E**). Since cabozantinib reduced *Tas1r2^+^* cells with a commensurate increase in GUST^+^ bitter/umami cells (see **Figure 2**), we hypothesized that TKI treatment would shift the fate of newly generated type II TBCs from sweet to bitter/umami fate. Consistent with this prediction, cabozantinib significantly increased the percentage of newly differentiated type II cells (Shh-Tomato^+^/PLCβ2^+^) that were GUST^+^ bitter/umami cells and significantly decreased the percentage that were *Tas1r2^+^*/ GUST^-^ sweet cells (**Figure 4F-I**). In sum, cabozantinib does not affect differentiation of type II cells in general but rather prevents differentiation of new *Tas1r2*^+^/GUST^-^ sweet cells, instead biasing new type II cells toward a GUST^+^ bitter/umami fate. Finally, these results explain why the impact of cabozantinib was greater with prolonged dosing (see **Figure 2**), as more TBCs were replaced over time with altered proportions of newly differentiated type II TBC subtypes.

### Cabozantinib impacts taste behavioral preferences of mice

Because sweet TBCs in the CVP were diminished by TKI treatment, we next asked whether sweet taste function was affected. To test this, we performed two behavioral assays: (1) a 48hr two-bottle taste preference test, and (2) a 30 min brief-access taste test. Mice were treated with vehicle or cabozantinib and tested behaviorally during the fourth week (**Figure 5A**). Mice prefer sweet, and when presented with a choice will consume/lick more sweet solution than water when sweet is detected above threshold (Reed and Knaapila, 2010). In two-bottle tests, neither vehicle- nor cabozantinib-treated mice preferred sweet solution at low concentrations of the non-nutritive sweetener SC45647 (Nofre et al., 1990). However, control mice strongly preferred 100 µM SC45647, while TKI-treated mice had no preference for sweet and drank equal volumes of sweetener and water (**Figure 5B**). Importantly, the total volume of solution consumed was not altered by drug treatment, although control mice drank significantly more 100 µM SC45647, likely driven by strong preference for the sweet solution (**Figure S7A**).

**Figure 5:**
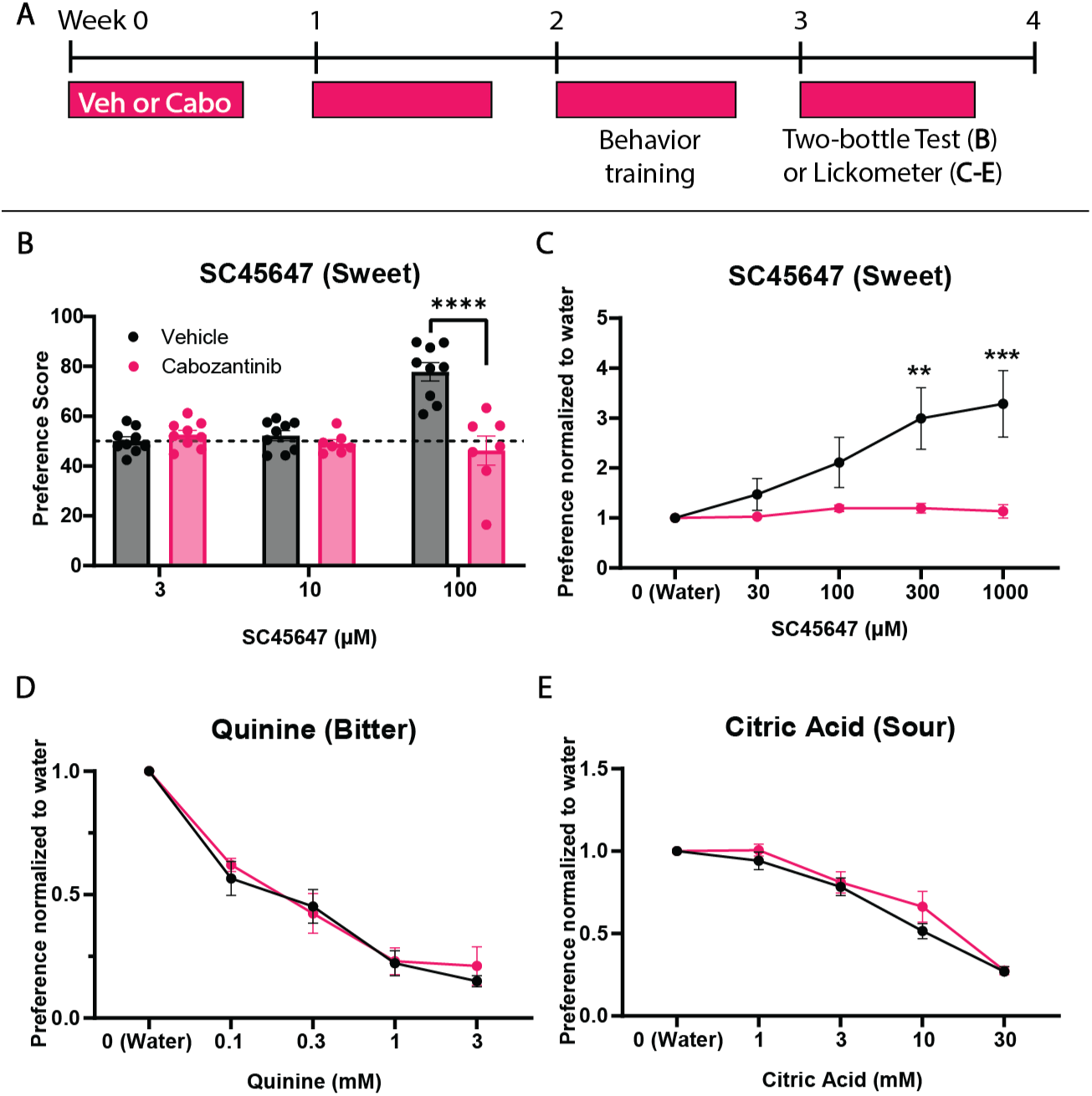
Sweet taste preference is diminished by cabozantinib. (**A**) Mice treated with vehicle or cabozantinib 5 days per week for 4 weeks underwent behavioral assay training in week 3, and taste behavior testing in week four. (**B**) In a two-bottle taste test, only control mice preferred sweet solution at high SC45647 concentration (N= 9 control and 7 TKI mice). Preference score = volume of SC45647 consumed / total volume of water and SC45647 consumed. Dotted line at 50 = no preference. (**C**) Brief access lickometer testing shows control mice prefer SC45647 at progressively higher concentrations while TKI-treated mice have no preference for SC45647 over water (N= 9 control and 9 TKI mice). Lickometer responses for quinine (**D**) and citric acid (**E**) were unaltered by TKI treatment (N= 9 control and 9 TKI mice). Preference = average licks for tastant / average licks for water. All data were analyzed using Two-way ANOVA with Sidak’s multiple comparisons test. Mean +/- SEM (** p≤0.01, *** p≤0.001, **** p≤0.0001).

A brief-access test with Davis Rig lickometers (Smith, 2001) was also used to assess taste behavior. This apparatus allows rapid testing of multiple concentrations of tastant in a 30-minute period. Moreover, limited access tests avoid post-ingestive effects that can influence taste behavior (Gaillard and Stratford, 2016). As with two-bottle testing, cabozantinib did not alter important metrics of drinking behavior (**Figure S7B-G**). Similarly, control mice in lickometer assays strongly preferred SC45647, while TKI-treated mice showed no preference for the non-nutritive sweetener (**Figure 5C**). Both two-bottle and lickometer tests reveal that cabozantinib decreases behavioral preference for sweet, consistent with the histological finding of reduced sweet TBCs in CVP taste buds.

Since cabozantinib increased bitter/umami type II cells, we also tested whether TKI treatment impacted bitter taste function. As bitter is aversive to mice, we hypothesized that the drug-induced increase in GUST^+^ cells would lower the threshold for avoidance of bitter solution. However, both vehicle- and cabozantinib-treated mice had comparable behavioral avoidance of quinine (**Figure 5D**). We also tested sour taste, which is mediated in part by type III TBCs (Liman et al., 2014). Avoidance of citric acid was not affected by drug treatment, consistent with our finding that cabozantinib did not impact type III cells (**Figure 5E**). Thus, cabozantinib did not affect taste preference for either bitter or sour but selectively blunted behavioral responses to sweet.

### In anterior FFP, cabozantinib does not affect production of sweet TBCs but instead reduces sodium-sensing TBCs

Although all taste qualities, including sweet, are detected by FFP and CVP taste buds, the FFP taste field is more sensitive to sweet tastants and thus we predicted that sweet type II TBCs in FFP would also be diminished by TKI treatment. In FFP, PLCβ2^+^ TBCs comprise 4 functional subtypes: sweet-, bitter-, umami-, and sodium-sensing. Unlike in the CVP, where *Tas1r2* and GUST are largely expressed by separate PLCβ2^+^ cell populations (sweet versus bitter/umami, respectively), almost all *Tas1r2*^+^ sweet cells in FFP co-express GUST, as do most bitter and umami type II cells (Kim et al., 2003, Tomonari et al., 2011). In contrast, specialized sodium-sensing TBCs express PLCβ2 and have strong developmental, molecular and morphological similarities with type II cells, but lack GUST expression (Ohmoto et al., 2020, Chandrashekar et al., 2010). Here, we considered all 4 functional TBC types as type II subtypes. We thus assessed if type II TBC subtype proportions were altered in FFP after 4 weeks of TKI dosing, and specifically if *Tas1r2*^+^/GUST^+^ sweet TBCs were reduced. Unexpectedly in FFP, neither *Tas1r2*^+^/GUST^+^ sweet cells nor GUST^+^ bitter/umami cells were impacted by drug treatment (**Figure S8I-J**). Instead, cabozantinib treatment resulted in fewer PLCβ2^+^/GUST^-^ sodium-sensing cells (**Figure S8E-H**).

We also assessed whether cabozantinib induced cell death or transdifferentiation of type II TBCs in FFP (see **Figure 3**). As in the CVP, cabozantinib had no impact on TBC survival as the number of *Kit*-YFP^+^ cells was unchanged (**Figure S9A-B**). Interestingly, while *Kit*-YFP primarily labels sweet type II cells in the CVP, *Kit*-YFP lineage traced into all three type II TBC subtypes at similar proportions in FFP; ∼60% of YFP^+^ cells were GUST^+^ (30% were Tas1r2^+^ sweet cells, meaning the other ∼30% were likely bitter/umami cells) and ∼40% were GUST^-^ sodium-sensing cells (**Figure S9C-H**). Regardless, cabozantinib did not change the percentage of *Kit*-YFP^+^ cells expressing PLCβ2, GUST or *Tas1r2* and thus did not induce transdifferentiation in FFP taste buds (**Figure S9C-H**).

Finally, we assessed the fate of TBCs newly differentiated from *Shh*^+^ precursor cells in FFP taste buds (see **Figure 4A**). As in the CVP, drug treatment did not affect differentiation of *Shh*-Tomato^+^ cells nor the percentage of *Shh*-Tomato^+^ cells differentiating as PLCβ2^+^ type II TBCs (**Figure S10A-D**). Instead, cabozantinib significantly increased the percentage of *Shh*- Tomato^+^/PLCβ2^+^ cells that were GUST^+^ and decreased the percentage that were GUST^-^ (sodium-sensing) (**Figure S10E-F**). Since differentiation of *Tas1r2*^+^/GUST^+^ sweet cells was unchanged (**Figure S10G-H**), the increase in *Shh*-Tomato^+^/GUST^+^ TBCs likely reflects an increase in differentiated bitter/umami type II cells at the expense of GUST^-^ sodium-sensing TBCs. Thus, cabozantinib alters the fate of newly generated type II TBCs, albeit different subtypes, in both CVP and FFP taste buds.

### Conditional deletion of *Kit* phenocopies TKI treatment in both CVP and FFP taste buds

Three lines of evidence suggested that TKI inhibition of the off-target RTK KIT may underlie shifts in type II TBC differentiation. In addition to intended targets VEGFRs and PDGFRβ, KIT is inhibited by all three TKIs tested here (Klaeger et al., 2017, Karaman et al., 2008, Davis et al., 2011). Further, mice treated with cabozantinib developed fur depigmentation (**Figure S11**), a physiological readout of KIT inhibition in mice and humans (Moss et al., 2003). Finally, *Kit* is expressed in differentiated sweet type II cells in the CVP and lineage traces into all type II cell subtypes in both taste fields, suggesting the type II cell lineage may be directly sensitive to KIT inhibition (see **Figures 3, S6, S9**). The POU2F3 transcription factor is expressed in all type II cells and is required for their differentiation (Matsumoto et al., 2011). Thus, we combined the *Pou2F3^CreER^* allele (Mcginty et al., 2020) with a novel floxed *Kit* allele (*Kit10*) wherein LoxP sites flank exon 10 that encodes the transmembrane domain of KIT protein (Lara et al. 2025, in revision). To delete *Kit* in the type II TBC lineage, *Pou2F3^CreER/+^;Kit ^fl/fl^* mice were dosed with tamoxifen or corn oil every other day for 2 weeks (**Figure S12A**).

Cre induction abolished KIT expression in CVP epithelium, indicating efficient excision of *Kit* (**Figure S12B-C**). *Kit* deletion did not affect numbers of type II or type III cells (**Figure S12D-G**), but as with drug treatment, significantly fewer *Tas1r2*^+^ sweet cells were detected (**Figure S12H-I**). Two weeks after *Kit* deletion, however, GUST^+^ cell number was unchanged (**Figure S12J-K**). We next tested if longer term *Kit* knockout would fully recapitulate the fate shift in type II TBC subtypes observed after 4 weeks of drug (see **Figure 2**). *Pou2F3^CreER/+^;Kit^fl/fl^*mice were fed tamoxifen chow for 4 weeks (**Figure 6A**). As tamoxifen chow can cause weight loss and other side effects (Halpage et al., 2024), control *Kit ^fl/fl^* mice lacking *Pou2F3^CreER^* were also fed tamoxifen chow. Under this protocol, KIT expression was efficiently ablated in CVP epithelium (**Figure 6B-C**). Consistent with 4 weeks of cabozantinib treatment, the number of type II and type III cells did not significantly change with prolonged *Kit* KO (**Figure 6D-G**), while *Tas1r2*^+^ sweet cells were significantly decreased (**Figure 6H-I**) with a commensurate, and significant, increase in GUST^+^ bitter/umami type II cells (**Figure 6J-K**). Thus, both cabozantinib and *Kit* KO induce a shift in type II TBC subtype fate after 4 weeks (**Figure 6L**).

**Figure 6:**
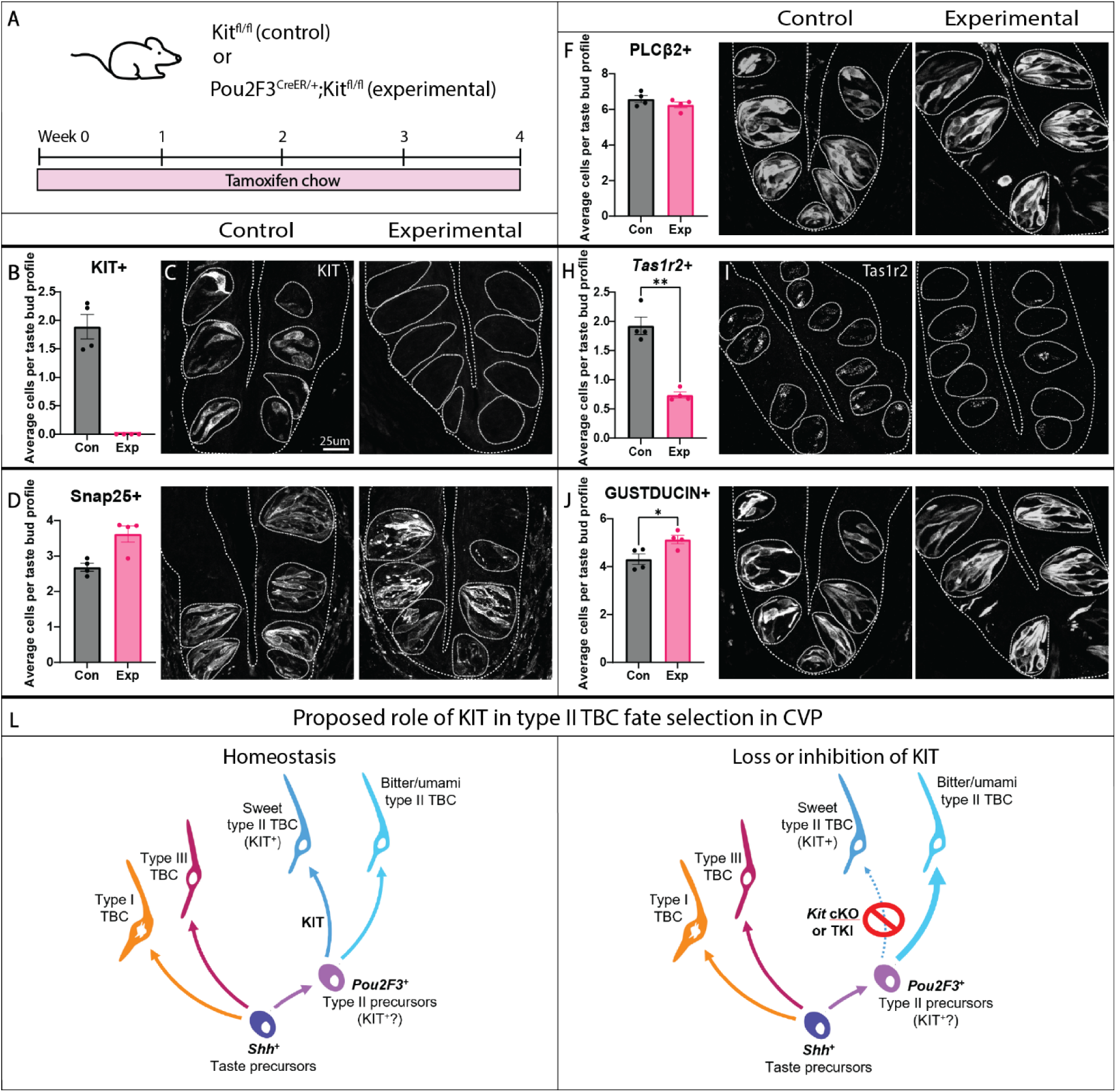
Conditional *Kit* knockout in the type II TBC lineage shifts type II TBC subtypes in CVP taste buds. (**A**) *Kit ^fl/fl^* (control) and *Pou2F3^CreER/+^;Kit ^fl/fl^* (experimental) mice were fed tamoxifen chow for 4 weeks. (**B-K**) Quantification of stained cells per taste bud profile with corresponding representative images for KIT-IF (**B-C**), SNAP25-IF (**D-E**), PLCβ2-IF (**F-G)**, *Tas1r2* HCR *in situ* hybridization (**H-I**) and GUSTDUCIN-IF (**J-K**). Representative images are compressed confocal z-stacks. Coarse dashed lines delineate basement membrane and apical epithelial surface, fine dashed lines encircle individual taste buds. Scale bar in **C** applies to **E**, **G**, **I** and **K**. In all histograms, each dot represents the average TBC tally from each mouse (N=4 per condition, ∼80 taste buds/mouse). Unpaired t-test performed on all quantifications. Mean+/-SEM (**** p≤0.0001). (**L**) Proposed model of *Kit* function in type II TBC lineage, where *Pou2f3^+^*precursor cells gives rise to all type II TBC subtypes, but those destined for a sweet cell fate upregulate *Kit*. Additionally, our pharmacological and genetic data support a model where KIT function is required for sweet type II cell fate in CVP. While different in details, we also conjecture a comparable model for FFP taste buds, but instead KIT function in *Pou2f3^+^*type II precursors is required for salt sensing type II cell fate (see **Figure S13**).

In FFP taste buds, POU2F3 is expressed by all PLCβ2^+^ cells, including sweet, bitter, and umami type II cells, as well as by PLCβ2^+^ sodium-sensing taste cells (Ohmoto et al., 2020). As in CVP, *Kit* KO in *Pou2F3*^+^ cells efficiently ablated *Kit* in FFP taste buds after two weeks of tamoxifen gavage (**Figure S13A-B**). As with drug treatment, conditional deletion of *Kit* did not affect the number of type II or type III cells (**Figure S13C-D, G-H**), nor were *Tas1r2*^+^ sweet cells affected (**Figure S13E-F**). Rather, consistent with the fate shift induced by cabozantinib in FFP, GUST^-^ sodium cells were decreased and GUST^+^ bitter/umami cells were increased by *Kit* deletion (**Figure S13I-L**). Thus, inducible *Kit* knockout phenocopies the effects of cabozantinib treatment in both FFP (**Figure S13M**) and CVP, suggesting TKIs disrupt type II TBC fate selection through KIT inhibition.

## Discussion

Many cancer drugs have the unintended side effect of perturbing taste. Since turnover of TBCs relies upon proliferating progenitors, therapies targeting proliferating cells are especially harmful to the taste system. For example, targeted head and neck irradiation, as well as anti-mitotic chemotherapies, are cytotoxic to taste progenitors and disrupt TBC renewal (Jewkes et al., 2018, Gaillard et al., 2019, Mukherjee et al., 2017, Ren et al., 2023). However, we find that the three anti-angiogenic TKIs tested here do not affect progenitor proliferation or survival when tested in lingual organoid cultures. TBC renewal can also be grossly disrupted by drugs that inhibit taste cell specification. Targeted drugs that inhibit SHH signaling, which is required for specification of new TBCs, prevent the specification of new taste cells and thus lead to progressive taste bud degeneration as older cells are lost and not replaced (Castillo et al., 2014, Castillo-Azofeifa et al., 2017, Liu et al., 2013, Kumari et al., 2015, Kumari et al., 2018, Yang et al., 2015, Lu et al., 2018). TKI treatment, however, did not cause large-scale disruption of TBC specification *in vivo* or *in vitro*. Instead, we find that differentiation of type II TBC subtypes specifically was dysregulated by anti-angiogenic drugs. In the CVP, cabozantinib decreased sweet type II cells and increased bitter/umami type II cells, while in FFP, sweet cells were not affected. Instead, cabozantinib led to reduced sodium cells with a commensurate increase in bitter/umami cells. Although multiple type II subtypes were altered by drug treatment, surprisingly only sweet taste behavior was impacted. In the CVP and FFP, these shifts in cell fate were recapitulated by deletion of *Kit* in the type II TBC lineage, suggesting that the impact of TKIs on type II TBC fate is due to KIT inhibition.

### Lineage dynamics of taste renewal

*Shh*^+^ taste precursor cells differentiate into all TBC types (Miura et al., 2014). However, it is not known whether these cells differentiate directly into TBCs or if further intermediate cell states exist wherein additional lineage decisions are made. *Ascl1* is expressed in type III TBCs and a subset of *Shh*^+^ precursors and is required for type III cell production (Seta et al., 2006, Seta et al., 2011). Lineage tracing of *Ascl1^+^* cells primarily labels type III cells and a small number of type II cells (Matsuyama et al., 2023, Hsu et al., 2021), suggesting *Shh^+^* precursors co-expressing *Ascl1* are capable of producing both cell types but are biased toward type III fate. In contrast, *Pou2F3* is expressed by some precursor cells and all mature type II TBCs and is necessary for type II cell production; *Pou2F3* knockout mice lose type II cells with a corresponding increase in type III cells (Matsumoto et al., 2011). Altogether, these studies suggest a model where *Shh^+^* precursor cells become progressively lineage restricted to type III and type II cell fate by *Ascl1* and *Pou2F3*, respectively. However, where in the taste lineage cells acquire type II TBC subtype fates is unknown. Since knocking out *Kit* in *Pou2f3*^+^ cells, as well as cabozantinib treatment, had no effect on differentiation of type II cells broadly but rather shifted the fate of type II TBC subtypes, we propose that *Shh^+^* descendent cells acquire type II subtype fate in POU2F3^+^ lineage-restricted type II cell precursors. Since FFP sodium cells also depend on POU2F3 for differentiation (Ohmoto et al., 2020), and since cabozantinib and *Kit* knockout affected the differentiation of sodium cells, our data support published findings that sodium cells and type II cells also arise from a common POU2F3^+^ precursor population in FFP taste buds (Ohmoto et al., 2020).

In both CVP and FFP, we find that *Kit* lineage tracing labels all type II TBC subtypes, albeit to different degrees, whereas KIT is not expressed in all differentiated type II cells (Choo and Dando, 2021, Ki et al., 2025). Together these findings suggest that KIT is expressed in POU2F3^+^ type II precursor cells. Additionally, conditional *Kit* knockout in *Pou2F3^+^* cells revealed that KIT is required for production of the correct proportions of type II TBC subtypes. Thus, we hypothesize that KIT signaling governs fate selection of differentiating type II TBCs. Interestingly, a recent report found that the KIT inhibitor imatinib had no effect on normal taste homeostasis (Ki et al., 2025). It is important to note that imatinib has a much shorter half-life than cabozantinib (∼20 hours versus ∼120 hours, respectively) (Peng et al., 2005, Lacy et al., 2017), and mice were treated daily for only 12 days. Since KIT inhibition likely affects type II cells as they gradually turnover, and since sweet cell reduction was observed after 14 days of either cabozantinib treatment or inducible *Kit* knockout, imatinib may not have been present at high enough levels and/or for long enough for the effects on type II cell differentiation to become apparent.

KIT function is required for differentiation of multiple cell types in other systems, including T-cells (Krishnamoorthy et al., 2008), cardiomyocytes (Li et al., 2008), spermatogonia (Nasimi et al., 2021), and melanocytes (Liao et al., 2017). Importantly, a complementary paper using *Pou2F3^CreER/+^;Kit ^fl/fl^* mice identified KIT as a regulator of intestinal tuft cell hyperplasia following helminth infection (Lara et al. 2025, in revision). In this context, KIT is required for the proliferation and/or differentiation of tuft cells during a type II immune response. Tuft cells, like type II TBCs, require POU2F3 for their generation (Gerbe et al., 2016), suggesting a shared requirement of KIT signaling in regulating the differentiation of POU2F3-dependent cells in multiple epithelial systems.

### Mechanisms of behavioral taste alterations

In rodents, it is well established that FFP taste buds in the anterior tongue are more highly tuned to sweet than posterior CVP taste buds, while posterior taste buds are more sensitive to bitter (Ninomiya et al., 1993, Shingai and Beidler, 1985). Thus, we were surprised when cabozantinib had no impact on sweet cell differentiation in FFP despite causing a loss of behavioral preference for sweet. Instead, sweet cell loss was evident only in CVP taste buds. These discordant changes in different taste papillae may contribute to TKI-induced dysgeusia, as incongruent input from anterior and posterior taste fields may change how taste stimuli are perceived (as proposed by (Gaillard et al., 2017)). Sweet taste behavioral responses may therefore depend on balanced sweet cell renewal in FFP and CVP, as normal sweet cell number in FFP did not appear to compensate for reduced sweet TBCs in the CVP. Additionally, it has been proposed that FFP taste buds mediate fine taste discrimination while CVP taste buds regulate a binary accept/reject response once substances reach the posterior oral cavity (Tomchik et al., 2007, Travers et al., 1987, St. John and Spector, 1998). Sweet cells in the CVP could be important to signal tastant acceptance, which may be required to drive appetitive behavior. Thus, a reduction in CVP sweet cells may lower tastant acceptance and decrease the appetitive quality of sweet.

We were also surprised that small reductions in sweet cell number resulted in such large effects on sweet taste behavior. KIT may therefore have additional functions in taste homeostasis. For example, KIT has known roles within the nervous system. *Kit* mutant mice display learning deficits and decreased synaptic potentiation in the hippocampus (Katafuchi et al., 2000). KIT signaling is also known to modulate synaptic activity between cerebellar Purkinje cells expressing KIT ligand and neighboring KIT^+^ interneurons. (Zaman et al., 2024). KIT may therefore function to modulate signaling between KIT^+^ taste cells and gustatory neurons. In support of this, transcriptome profiling of gustatory neurons in the geniculate ganglion, which innervate FFP taste buds, revealed expression of both *Kit* and KIT ligand (*Kitl*), suggesting KIT function may be important for proper transmission of taste information to the brain (Zhang et al., 2019, Anderson and Larson, 2020). Additionally, recent findings suggest that KIT function confers resistance to sweet cell loss following nerve injury, and that these residual sweet cells may promote re-innervation of taste buds by regenerating nerve fibers (Ki et al., 2025). Together, these studies suggest that the loss of sweet preference in cabozantinib-treated mice could be due to effects on sweet-sensing gustatory neurons in addition to the effects on type II TBC differentiation.

On the other hand, behavioral preferences for bitter and sour tastants were unaffected by TKI treatment. Since GUST expression is associated with bitter cells in both CVP and FFP taste buds (Kim et al., 2003, Tomonari et al., 2011), we expected the increase in GUST^+^ type II cells to cause hypersensitivity to bitter. However, behavioral sensitivity to bitter compounds may already be maximal. Many plants, fungi, and animals produce toxic or poisonous substances to deter predation; these are almost always bitter, and thus the taste system may have evolved to detect and allow avoidance of bitter substances at low concentrations (Reed and Knaapila, 2010). If bitter taste sensitivity is already high, small increases in bitter cells would have little functional consequence. Overall, our data suggest that loss of sweet taste function is a primary cause of TKI-induced dysgeusia. Intriguingly, sweet taste was reportedly the most commonly affected taste modality in patients treated with TKIs for gastrointestinal stromal tumors, while salt taste was the second most commonly affected taste modality (Van Elst et al., 2022).

### Limitations and future directions

While TKI treatment decreased sodium-sensing type II cells in FFP taste buds, we did not test salt behavior in TKI-treated mice. FFP sodium TBCs mediate detection of low, appetitive concentrations of salt (Ohmoto et al., 2020). Practically, motivating mice to prefer such low concentrations requires dietary salt depletion that may not be well tolerated by TKI-treated mice (Chandrashekar et al., 2010). Additionally, while the taste system of mice is broadly similar to that of humans (Tizzano et al., 2015), we can only speculate that humans experience similar cell type shifts and changes in taste preferences. This limitation could be addressed in the future by collecting taste bud biopsies from TKI-treated patients to investigate whether cellular changes are similar to those observed in mice, and by performing psychophysical taste testing for individual tastants (Doty, 2019) in patients experiencing dysgeusia.

In closing, given the importance of sweet cells in the maintenance of sweet taste preference, and given KIT’s role in regulating sweet cell renewal in CVP taste buds, we propose that KIT inhibition contributes to TKI-induced dysgeusia. In addition to the three TKIs tested here, KIT is an off-target RTK of seven anti-angiogenic TKIs currently used in clinical practice (Klaeger et al., 2017, Chang et al., 2022, Sánchez-Gastaldo et al., 2017). Thus, KIT inhibition may underlie dysgeusia in this large class of targeted cancer therapies. For drugs used to treat mRCC, KIT is not an intended target, as therapeutic efficacy relies primarily on VEGFR and PDGFRβ inhibition (Sánchez-Gastaldo et al., 2017). Developing alternatives that maintain inhibition of these targets without inhibiting KIT could alleviate dysgeusia while maintaining efficacy in treating mRCC. There are also cancers in which oncogenic KIT mutations drive tumorigenesis, such as gastrointestinal stromal tumors, melanomas, small cell lung cancer and many leukemias (Lennartsson and Rönnstrand, 2012). Unfortunately, treating these cancers requires KIT inhibition and therefore dysgeusia is likely unavoidable. Developing targeted therapies that specifically inhibit mutated KIT without inhibiting wild-type KIT may mitigate dysgeusia in these cases. By establishing KIT as a regulator of type II TBC subtype renewal and showing that *Kit* knockout underpins the response to TKI treatment, we have provided a promising candidate for future mitigation strategies.

## Materials and Methods

### Animals

Commercially available mice were obtained from the Jackson Laboratory (*Lgr5^EGFP-IRES-CreERT2^*, 008875; *Shh^CReERT2^,* 005623*; Rosa26^TdTomato^,* 007909*; Rosa26^YFP^,* 006148; *Pou2F3^CreERT2-IRES-eGFP^*, 037511). *Kit^CreER/+^;Rosa26^YFP/YFP^* mice were a gift from Stephen W. Santoro, University of Colorado Anschutz Medical Campus, Aurora CO (originally reported (Klein et al., 2013)). *Pou2F3 ^CreERT2-IRES-eGFP/+^;Kit ^fl/fl^* mice were a gift from Jakob von Moltke, University of Washington, Seattle WA (Lara et al. 2025, in revision). Mice were maintained in an AAALAC-accredited facility in compliance with the Guide for Care and Use of Laboratory Animals, Animal Welfare Act, and Public Health Service Policy. Male and female adult mice between 8-20 weeks were used. Procedures were approved by the Institutional Animal Care and Use Committee at the University of Colorado Anschutz Medical Campus.

### Administration of drugs

<u>Cabozantinib malate</u> (XL 184, SelleckChem) was suspended in HPLC water at 6 mg/ml with 5 µl/ml 1N HCL (Fisher Chemical), sonicated for 10 min, heated at 37°C for 15 min with vortexing, and sonicated for another 10 min. Adult mice on mixed background were individually housed and dosed with 60 mg/kg cabozantinib or vehicle via oral gavage. Mice were weighed daily to monitor health.

<u>Tamoxifen gavage</u>: Tamoxifen (Sigma-Aldrich) was suspended in corn oil + 10% ETOH at 10 mg/ml. Adult mice were individually housed and gavaged with 50 mg/kg (*Shh^CreER/+^;Rosa^26tdTomato/+^*) or 100 mg/kg (*Kit^CreER/+^;Rosa26^YFP/YFP^* and *Pou2F3^CreER/+^;Kit ^fl/fl^*). Cages were changed daily for 7 days after final tamoxifen dose or until mice were harvested, whichever came first.

<u>Tamoxifen chow:</u> Control (*Kit ^fl/fl^*) and experimental (*Pou2F3^CreER/+^;Kit ^fl/fl^*) mice were individually housed and fed Tamoxifen chow (Inotiv, 500) ad libitum for 4 weeks. Mice were weighed daily for the first 10 days to monitor weight loss, then weighed three times per week for the remainder of the study once their weights recovered and stabilized.

### Behavioral Assays

<u>Two-bottle taste preference test</u>: Preference for SC45647 was tested in a 48 hr two-bottle behavioral assay (Gaillard and Stratford, 2016). Mice were individually housed in regular ventilated cages with *ad libitum* chow. Graduated bottles were created from cut plastic serological pipettes fitted with a sipper tube on one end and a rubber stopper on the other. Mice were trained for 4 days; first with deionized water in one of 2 bottles, and positions switched after 24 hrs. In the second 2 days, mice had access to water in both bottles. To test sweet taste preference, mice had one bottle of water and one bottle of SC45647 (3 µM in deionized water) for 48 hrs; the volume consumed was measured at 24 hrs, bottles refilled as necessary, and left-right position switched to control for side preference. Sweet preference ratio was calculated by dividing the volume of SC45647 consumed by the sum of the volume of water and SC45647 consumed over 48 hrs. This was repeated for 10 µM and 100 µM SC45647.

<u>Brief Access Test (Lickometer)</u>: Three Davis Rig MS-160 lickometers (DiLog Instruments, Inc., Tallahassee, FL, USA) were used to test taste preference of vehicle- and cabozantinib-treated mice. Detailed protocol and operating procedures have been described fully (Gaillard and Stratford, 2016). During the last 2 weeks of drug or vehicle treatment, mice were trained and tested in one of three lickometers randomly assigned each day. Chamber areas were reduced by half to reduce exploration and distraction from surroundings. Training and test sessions were limited to 30 min. During brief-access testing a shutter opens to give mice access to a bottle of tastant for 5 sec (counted from first lick), the shutter closes for 7.5 sec, and the next bottle is moved into position via an automated mobile rack for the next 5 sec trial. Mice were deprived of water for 23.5 hrs prior to training or testing to motivate drinking. Mice were trained to drink water in the lickometer for 4 consecutive days: For 2 days, the shutter was open and water available for 30 min; then on days 3 and 4, two bottles of water were alternately presented for 5 sec each for 30 min. After 2 days of recovery, mice had a final day of training with two alternating bottles of water. Once trained, mice were tested for behavioral taste preference for 4 days. For each cabozantinib versus vehicle experiment, only 2 of a total of 3 tastants were assayed, each for 2 days. These included: Sweet - SC45647 (30, 100, 300 and 1000 μM), Bitter - quinine (0.1, 0.3, 1 and 3 mM) and Sour - citric acid (1, 3, 10 and 30 mM). All solutions were made in deionized water. Each test session consisted of up to eight trial blocks (each block comprised five presentations in random order - water and 4 concentrations of one tastant). For sweet SC45647, the first block was not included in our analysis as a thirst-induced high motivational state during the first block may drive maximal licking for all solutions (Gaillard and Stratford, 2016). Mouse thirst for all tastants was calculated from the total licks across the first block averaged across the two testing days. Results of behavior tests are expressed as the lick ratio (average licks to tastant/average licks to water).

### Tissue Preparation

Mice were euthanized by CO2 asphyxiation. Tongues were dissected from the lower jaw, incubated for 3 hrs at 4°C in 4% paraformaldehyde (PFA) in 0.1 M phosphate buffer (PB), and placed in sucrose (20% in 0.1 M PB) overnight at 4°C. Samples were embedded in OCT Compound (Tissue-Tek), 12 µm cryosections collected on SuperFrost Plus slides (Thermo Fisher Scientific) and stored at −80°C. For the CVP, 6 sets of 9 sections were collected, while for the FFP, 6 sets of 16 sections were collected.

### Immunofluorescence

<u>Tissue sections</u>: Sections were washed in 0.1 M PBS, incubated in blocking solution (BS: 5% normal goat or donkey serum, 1% bovine serum albumin, 0.3% Triton X-100 in 0.1 M PBS at pH 7.3) for 1.5 hrs RT followed by primary antibodies (Table S2) diluted in BS overnight at 4°C. Sections were rinsed in PBS + 0.1% Triton, incubated for 1 hr at RT in secondary antibodies in BS, followed by DAPI nuclear counterstaining and washes with 0.1 M PB (Invitrogen; 1:10,000). Slides were cover-slipped with ProLong Gold (Thermo Fisher Scientific).

<u>Organoids</u>: Organoids were generated from taste progenitor cells obtained via FACS of dissociated CVP epithelia from *Lgr5^EGFP-IRES-CreERT2^* mice following our published protocol (Shechtman et al., 2021). To harvest, organoids were incubated in Cell Recovery Solution (Corning) at 4°C, washed in 0.1 M PBS, fixed in 4% PFA and stored at 4°C in PBS with 1% BSA. For immunofluorescence, organoids were incubated in BS (2 hrs), then with primary antibodies (Table S2) in BS for 3 nights at 4°C, washed with PBS + 0.2% Triton and incubated with secondary antibodies in BS overnight at 4°C. Organoid nuclei were counterstained with DAPI, washed with 0.1 M PB and mounted on SuperFrost Plus slides in Fluoromount (Southern Biotech).

### Hybridization chain reaction RNA-fluorescence *in situ* hybridization

Molecular Instruments designed and produced probes against *Tas1r2* (NM_031873.1), *Kit* (NM_001122733.1) and *Trpm5* (NM_020277.2). Methods were adapted from the manufacturer’s protocol. Fixed frozen sections were incubated in 4% PFA for 10 min at RT, washed with 0.1X PBS and incubated in 2 µg/ml Proteinase K for 2 min (CVP) or 5 min (FFP) at RT. Sections were incubated in triethanolamine solution, 12 M HCl and acetic anhydride in DEPC-treated water for 10 min at RT, washed with 0.1X PBS, incubated in hybridization buffer for 45 min at 37°C, and then with 1.2 pmol (CVP) or 1.5 pmol (FFP) probe in hybridization buffer overnight at 37°C. Sections were washed with 75% wash buffer/25% 5× SSCT (20× SSC and 10% Tween20, ultrapure water), 50% wash buffer/50% 5× SSCT, 25% wash buffer/75% 5× SSCT, and 100% 5× SSCT, each for 20 min at 37°C, then with 100% 5× SSCT for 20 min at RT. Sections were incubated in amplification buffer for 1 hr, then in denatured hairpin solution (6 pmol hairpin 1 and 2 in amplification buffer) overnight. Slides were washed with 5× SSCT, nuclei counterstained with DAPI, washed with 0.1 M PB and cover-slipped with ProLong Gold.

### Organoid Derivation

Organoids were derived according to our published protocol (Shechtman et al., 2021). For each experiment, organoids were derived from CVP epithelium from 3-6 *Lgr5^EGFP-IRES-CreERT2^* mice aged 8-20 weeks. Briefly, collagenase (2 mg/ml) and dispase (5 mg/ml) in phosphate-buffered saline (PBS) were injected around the CVP, then the epithelium peeled and dissociated for 45 min in collagenase (2 mg/ml), dispase (5 mg/ml) and elastase (2 mg/ml) at 37°C. Cells were centrifuged (320 ***g*** at 4°C), the pellet resuspended in FACS buffer [1 mM EDTA, 25 mM HEPES (pH 7.0), 1% FBS, 1×Ca^2+^/Mg^2+^-free dPBS], passed through a 30 µm cell strainer, stained with DAPI (Thermo Fisher Scientific) and subjected to FACS on a MoFlo XDP100 (Cytomation). Debris and doublets were gated out via side-scatter (FSC and SSC-width, respectively), live (DAPI^neg^) cells were enriched, and the gating of GFP^+^ signal was determined relative to background fluorescence of epithelial cells from a wild-type control mouse processed in parallel (see (Shechtman et al., 2021) for gating strategy). Only *Lgr5*-GFP^+^ cells were collected. Sorted GFP^+^ cells were plated in 48-well plates at 200 cells/well in 15 µl Matrigel and grown in WENR+AS media to support growth (days 0-6) and WENR media to promote TBC differentiation (days 6-12). WENR: 50% WRN (WNT/RSPO/Noggin)-conditioned media (Miyoshi and Stappenbeck, 2013) plus 1× Glutamax (Gibco), 1× HEPES (Gibco), 1× penicillin-streptomycin (Gibco), 1× B27 supplement (Gibco), 1× gentamicin (Gibco), 1× primocin (InvivoGen), 25 ng/ml murine Noggin (Peprotech), 50 ng/ml murine EGF (Peprotech), 1 mM nicotinamide (Sigma), and 1 mM N-acetyl-L-cysteine (Sigma). WENR+AS: WENR plus 500 nM A83-01 (Sigma) and 0.4 µM SB202190 (R&D Systems). Y27632 (10 µm; Stemgent) was added on days 0-2 to promote cell survival.

<u>Drug treatment</u>: cabozantinib (SelleckChem; S4001), axitinib (SelleckChem; S1005), sunitinib (SelleckChem; S7781), and paclitaxel (SelleckChem; S1150) were dissolved in DMSO to a stock concentration of 50 µM, and diluted in organoid media (WENR+AS or WENR depending on the experiment) to a final concentration of 50 nM or 100 nM. For controls, the same volume of DMSO used for 100 nM drug was added to organoid media. Medium was changed every 2 days.

### Quantitative RT-PCR

Organoids were harvested as described (Shechtman et al., 2021). Briefly, plates were placed on ice for 30 min and organoids freed from Matrigel by scratching with a pipet tip. Organoid samples were centrifuged and resuspended in RLT buffer (Qiagen) with 1% β-mercaptoethanol. RNA was extracted using a RNeasy Micro Kit (Qiagen), quantified via Nanodrop (Thermo Fisher Scientific) and reverse transcribed with an iScript cDNA synthesis kit (Bio-Rad). Power SYBR Green PCR Master Mix (Applied Biosystems) was used for qPCR reactions on a StepOne Plus Real-Time PCR System (Applied Biosystems, Life Technologies). Relative gene expression was assessed using the ΔΔCT method (Livak and Schmittgen, 2001), with *Rpl19* as the housekeeping gene. Primers are listed in **Table S1**. Organoids from three wells of a 48-well plate were pooled per RNA sample and three samples were collected per experiment for RT-qPCR.

### EdU incubation, detection and quantification

Organoids were grown in 48-well plates as described above and on day 6, EdU (4 mM stock in 0.9% NaCl) was added to WENR+AS media at a final concentration of 10 µM for 30 min. Cultures were placed in fresh WENR+AS on ice and organoids harvested for immunofluorescence as above. A Click-it® EdU Alexa Fluor® 647 Imaging Kit (Invitrogen) was used to detect EdU signal using methods adapted from the manufacturer’s protocol. Organoids were washed with PBS +0.2% Triton, washed with PBS + 0.1% BSA, then incubated in Click-it® Plus reaction cocktail for 3 hrs at RT. Organoids were washed with PBS, nuclei counterstained with DAPI, washed with 0.1 M PB and mounted on SuperFrost Plus slides in Fluoromount (Southern Biotech). Using a custom app - OrganoidAnalyzer in MATLAB 2024a (Mathworks, Natick, MA) (https://github.com/salcedoe/OrganoidAnalysis) - we determined the ratio of EdU^+^ pixels to total DAPI^+^ pixels in each organoid. See supplemental materials for details.

This app allows the user to select and open a confocal image stack and visually inspect each stack by channel and optical section. The app also provides the controls to process and segment the stack, and capture and collate the data. The app uses the Bio-Formats MATLAB toolbox (Linkert et al., 2010) to open each image stack. Once a confocal z-stack is opened, each channel in the stack is processed separately. Image stacks are normalized to the maximum intensity found in the channel (using the **mat2gray** function). Stacks are then filtered using a median filter with the default neighborhood settings (**medfilt3**) and a gaussian filter using a 5 x 5 x 5 filter size (**imgaussfilt3**). To increase contrast and enhance separation between nuclei in an organoid, we create an image stack marker by applying an erosion morphological operation on the filtered stack (**imerode)** using a sphere-shaped structuring element with a radius of 3 voxels (**strel**). We then perform a morphological reconstruction of the filtered volume using the image stack marker we created in the previous step (**imreconstruct)**. To identify labeled nuclei, image stacks are segmented into labeled nuclei and background using a modified version of Otsu’s Method for thresholding. Specifically, a threshold value is calculated for each channel using **graythresh** and this value is multiplied by a set factor (i.e. 0.5 for the DAPI channel and 0.2 for the EdU channel). The same factors are used for all image stacks to ensure consistent segmentation. To create the binarized segmentation image stack, pixels with intensity values greater than the threshold value are labeled with a logical one to signify labeled cells, while pixels with intensity values below the threshold value are labeled with logical zeros to signify background. We created two separate segmentation stacks: one for the EdU-labeled cells and one for the DAPI-labeled cells. Each segmentation stack is visually inspected and compared to the original image stack to ensure that pixels designated as signal were in fact signal and not noise. The ratio of EdU^+^ nuclei to total DAPI^+^ nuclei was calculated by dividing the total number of logical ones in the EdU binarized image stack by the total number of logical ones in the DAPI binarized image stack. As such, we are in effect dividing the total volume of EdU^+^ nuclei by the total volume of all DAPI^+^ nuclei to calculate the ratio. This was necessary as our technique was unable to separate all individual nuclei in an organoid, especially the DAPI^+^ nuclei which are tightly clustered together.

### Cell-Titer Glo® 3D

Organoids were derived and cultured as above but in opaque-walled 96-well plates at 100 cells per well in 3 µl Matrigel. On day 6, plates were equilibrated to RT for 30 min, after which an equal volume of Cell-Titer Glo® 3D reagent (Promega) was added to cultures (150 µl WENR+AS with drugs or DMSO + 150 µl reagent in each well). Plates were mixed on an automatic plate shaker for 5 min to induce cell lysis. Luminescence was recorded after 25 min using a Synergy H1 microplate reader (BioTek). The average luminescence across 6 wells was calculated for each condition in each biological replicate.

### Image acquisition and analysis

<u>Immunofluorescence</u>: Tissue sections and organoids were imaged with a Leica TCS SP8 laser-scanning confocal microscope with LAS X software. *Z*-stacks of optical sections of CVP and FFP sections were acquired at 0.75 µm thickness, and organoids at 2 µm optical section thickness. For image acquisition, analysis, and cell counts using LAS X Office software, investigators were blinded to condition. For CVP, all taste buds in 5 tissue sections were counted. For FFP, all taste buds in 16 sections from the tongue tip were counted. Criteria for immunolabeled cell counts: (1) cell is immunomarker-positive; and (2) has a DAPI^+^ nucleus. Taste buds were identified during cell counting using either PLCβ2 staining (co-staining with GUST and YFP), KRT8 staining (co-staining with SNAP25 and KIT), or *Trpm5* HCR *in-situ* hybridization (with *Tas1r2* probe).

<u>Taste bud area</u>: Maximum z-stack projections of KRT8 immunofluorescence tissue sections were generated and imported into ImageJ software. A scalebar was used to set pixels/µm scale. Individual taste buds were outlined using the Freehand Selection tool to obtain the area of each taste bud profile.

### Statistical analyses

In experiments with two conditions, normally distributed data were analyzed by unpaired t-test, and Mann-Whitney U tests were used when data were not normally distributed. In experiments with more than two conditions, normally distributed data were analyzed by ordinary one- or two-way ANOVA with either Sidak’s or Tukey’s multiple comparisons post-hoc test. A Kruskal-Wallis comparison test was used when data were not normally distributed. All statistical analyses employed GraphPad Prism software. Data plotted throughout are depicted as Mean ± S.E.M. and significance was taken as *p*<0.05 with a confidence interval of 95%.

## Acknowledgments

The authors thank the University of Colorado Organoid and Tissue Modeling Shared Resource (OTMSR) for providing WRN conditioned media and technical assistance for organoid experiments. The authors thank Dmitry Baturin and Lester Acosta of the University of Colorado Cancer Center Flow Cytometry Shared Resource for FACS, Trevor J. Isner and Amanda Stenzel for technical assistance with mouse and organoid experiments, and Thomas E. Finger, Katherine Fantauzzo, Heide Ford, Stephen Santoro, Santos Franco, Trevor J. Isner and Ian J. Purvis for helpful discussions and for critically reading the manuscript. Finally, we heartily thank Robin F. Krimm for sharing unpublished data.

## Funding

National Institutes of Health T32GM141742 (CMP)

National Institutes of Health F31DC020634-01 (CMP)

National Institutes of Health R01DC021865 (LAB)

National Institutes of Health R01DC018489 (LAB)

National Institutes of Health R01DC012383 (LAB)

National Institutes of Health R21CA236480 (LAB)

National Institutes of Health R01AI167923 (JVM)

National Institutes of Health P30CA046934 (CU OTMSR)

National Institutes of Health P30DK116073 (CU OTMSR)

National Institutes of Health P30CA046934 (CU Cancer Center)

## Author contributions

Project conceptualization: CMP, LAB, PJD, ETL

Organoid experiments: CMP, JKS

MATLAB coding and custom app creation: ES

Mouse handling: CMP, JKS, CEW

Tissue processing: CMP, JKS, ASH

Imaging and cell counting: CMP, ASH

Behavior: CMP, CEW, ASH

Generation of novel KIT10 allele: HIL, JVM

Writing: CMP, ES, LAB

Figure creation: CMP, LAB

Editing: all authors

## Data and materials availability

All relevant data and details of resources can be found within the article and its supplementary information.

## Supplementary Material

**Figure S1.**
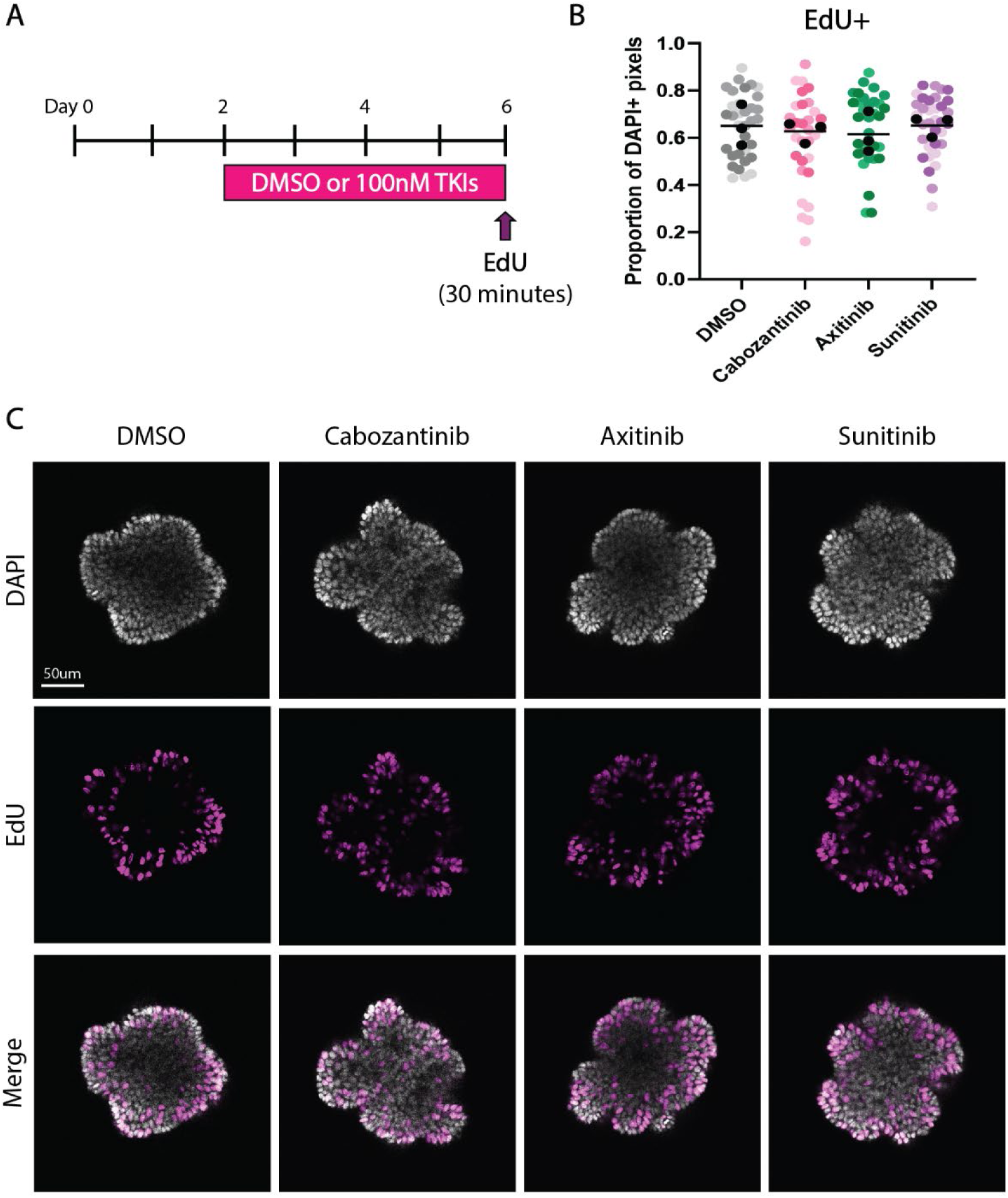
TKIs do not affect proliferation of taste progenitor cells in organoids. (**A**) Timeline of lingual organoid treatment with DMSO or 100 nM TKIs during the growth phase (day 2-6). EdU was added to the media for 30 minutes prior to harvest on day 6. (**B**) Drug treatment did not affect EdU labeling of organoids. Each colored dot represents the proportion of DAPI^+^ pixels labeled with EdU in a single organoid. Within a treatment condition, different shades represent the 3 biological replicates, and black dots represent averages of each replicate. Ordinary one-way ANOVA with Dunnett’s multiple comparisons test was used to compare the average values (black line) across conditions.

**Figure S2:**
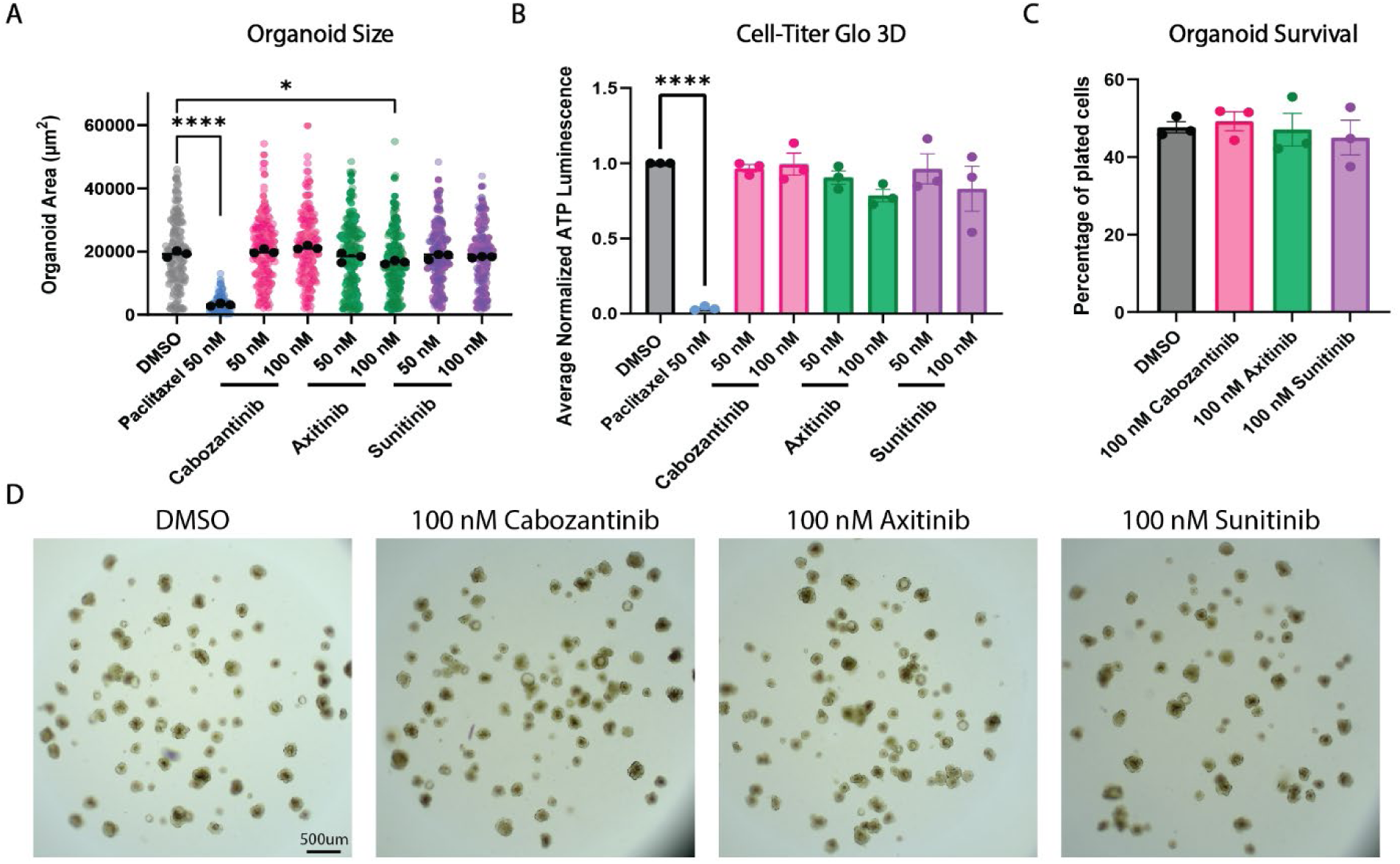
TKI treatment during the growth phase of culture does not affect growth or survival of lingual organoids. (**A**) Organoid area was not broadly affected by treatment with 50 or 100 nM TKIs (see Fig 1A) compared to negative control (DMSO), but was significantly reduced by paclitaxel (positive control). Each colored dot represents the area of one organoid. Within a treatment condition, different shades represent the 3 biological replicates, and black dots represent averages of each replicate. (**B**) Estimation of cell survival using Cell-Titer Glo^®^ 3D was unaltered by TKI treatment compared to DMSO control, but was significantly diminished by paclitaxel. Each dot represents the average luminescence across 6 pooled wells for each of 3 biological replicates. (**C**) Organoid survival did not differ with condition. The number of organoids in each well at day 6 was divided by the number of plated cells at day 0 (200 cells per well). Each dot represents the average of three wells for each of 3 biological replicates. Ordinary one-way ANOVA with Dunnett’s multiple comparisons test was performed on experimental averages in panel A and on all values in panels B and C. Mean +/- SEM (* p≤0.05, **** p≤0.0001). (**D**) Brightfield images of culture wells containing organoids treated with DMSO or 100 nM TKIs.

**Figure S3:**
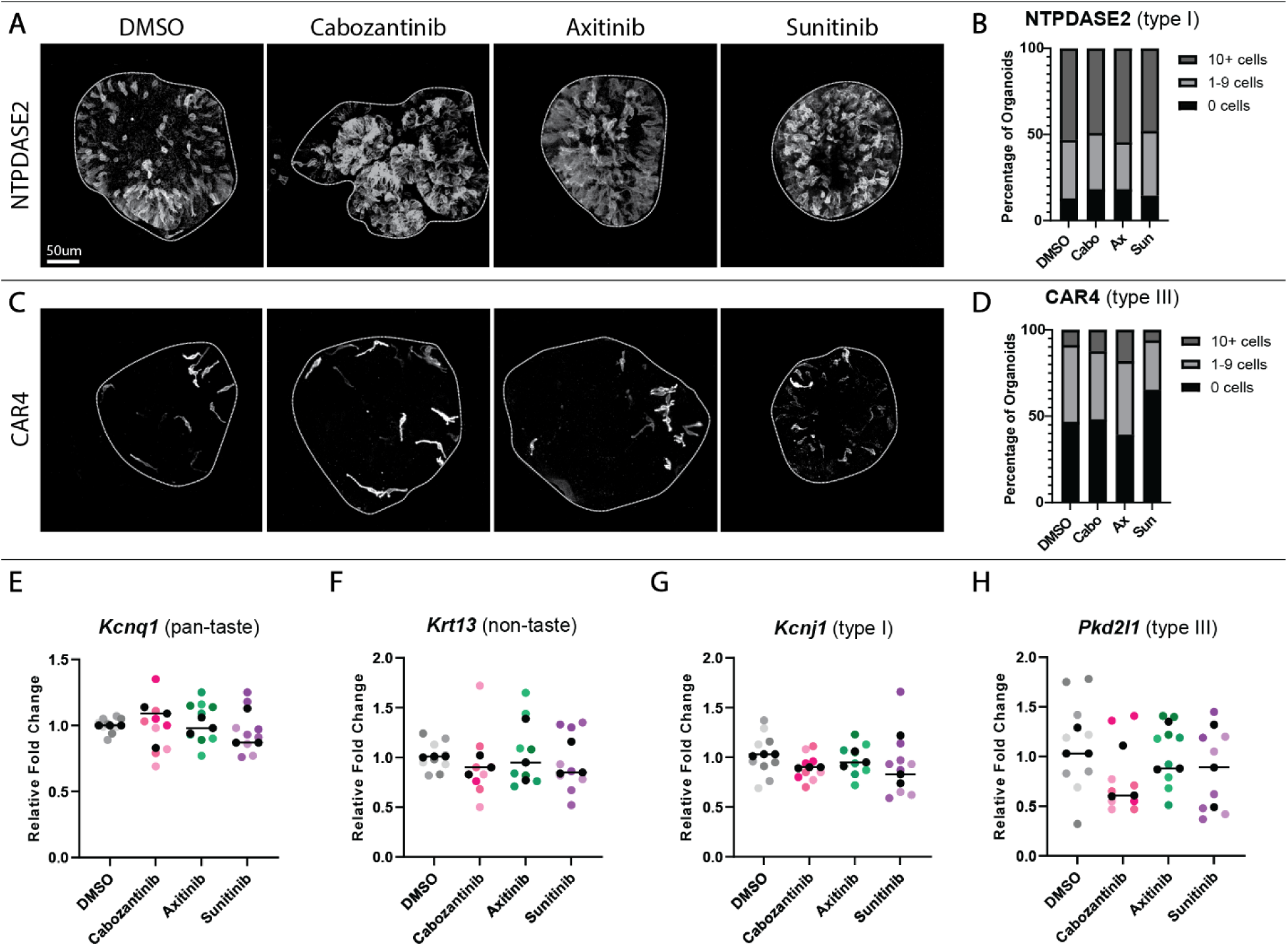
TKIs do not affect type I or type III TBCs or markers of taste vs non-taste epithelium. Compressed confocal z-stacks of control and TKI-treated organoids immunostained for markers of type I (NTPDase2) (**A**) and type III (CAR4) TBCs (**C**). Histograms showing percentages of organoids containing 0, 1-9, or ≥ 10 cells immunopositive for NTPDase2 (**B**) or CAR4 (**D)** reveal no change with drug treatment compared to controls. Total organoids obtained from 3 biological replicates: DMSO - 47; cabozantinib - 55; axitinib - 33; sunitinib - 48. Relative fold change in expression of pan-taste marker *Kcnq1* (**E**), non-taste marker *Krt13* (**F**), type I TBC marker *Kcnj1* (**G**) and type III TBC marker *Pkd2l1* (**H**) measured via RT-qPCR. Each colored dot represents an individual sample where organoids were pooled from three culture wells (∼240 organoids per sample). Within a treatment condition, different shades represent the 3 biological replicates, and black dots represent averages of each replicate. Ordinary one-way ANOVA with Dunnett’s multiple comparisons test was used to compare the average values (black line) for each marker across conditions.

**Figure S4:**
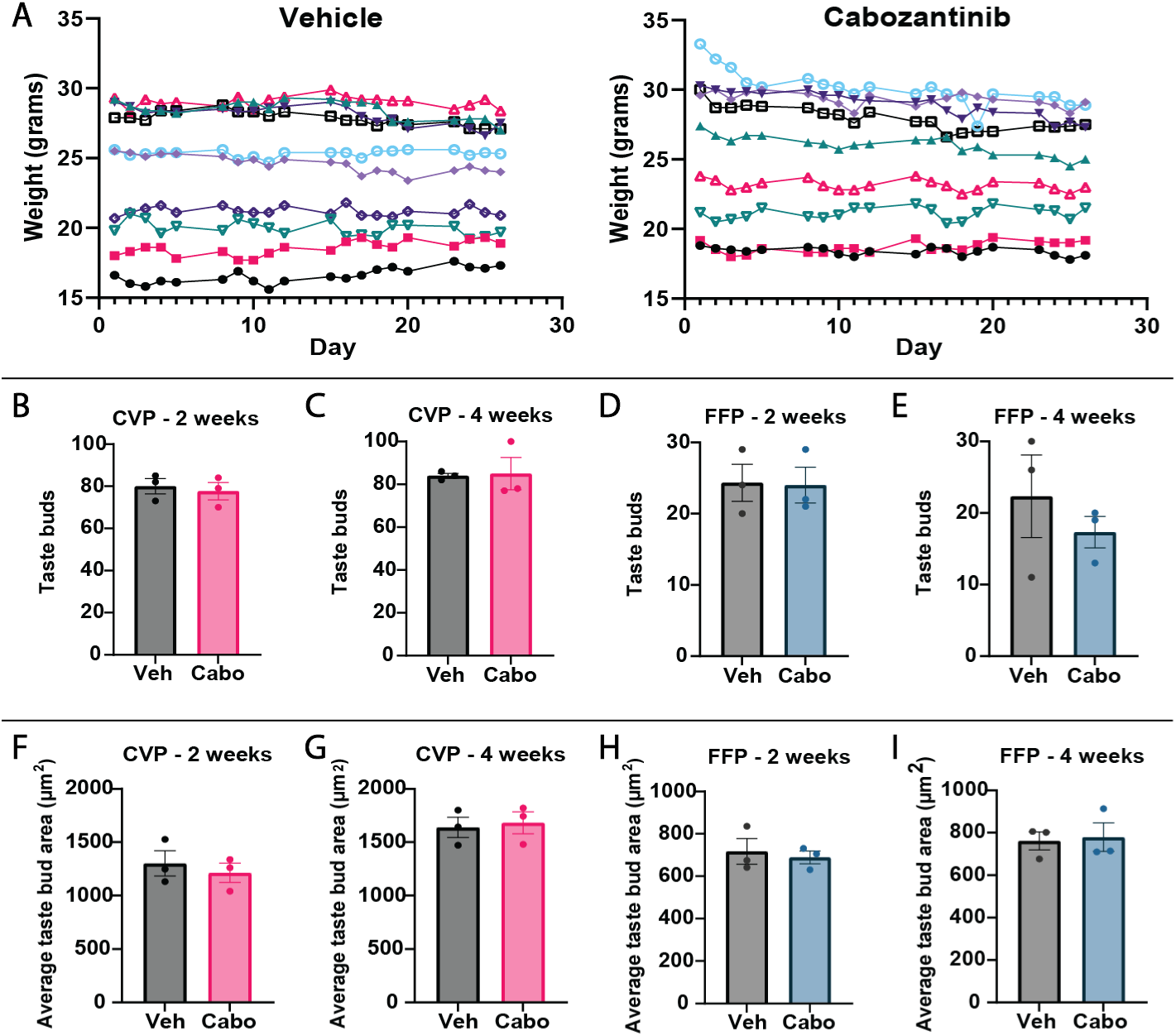
Cabozantinib does not affect mouse health or general taste bud homeostasis. (**A**) Weights of mice treated with vehicle or cabozantinib did not differ over 28 days. Weight was measured 5 days per week (symbols) and each colored line is an individual mouse (N=9 mice per condition). The number of CVP taste buds did not differ with treatment at 2 weeks (**B**) or 4 weeks (**C**). The number of FFP taste buds in the first 1.2 mm of the tongue was unaltered by cabozantinib treatment at 2 weeks (**D**) or 4 weeks (**E**). The average area of CVP (**F-G**) and FFP (**H-I**) taste bud profiles did not differ with treatment either at 2 weeks or 4 weeks (N=3 mice per condition). Unpaired t-test. Mean +/- SEM.

**Figure S5:**
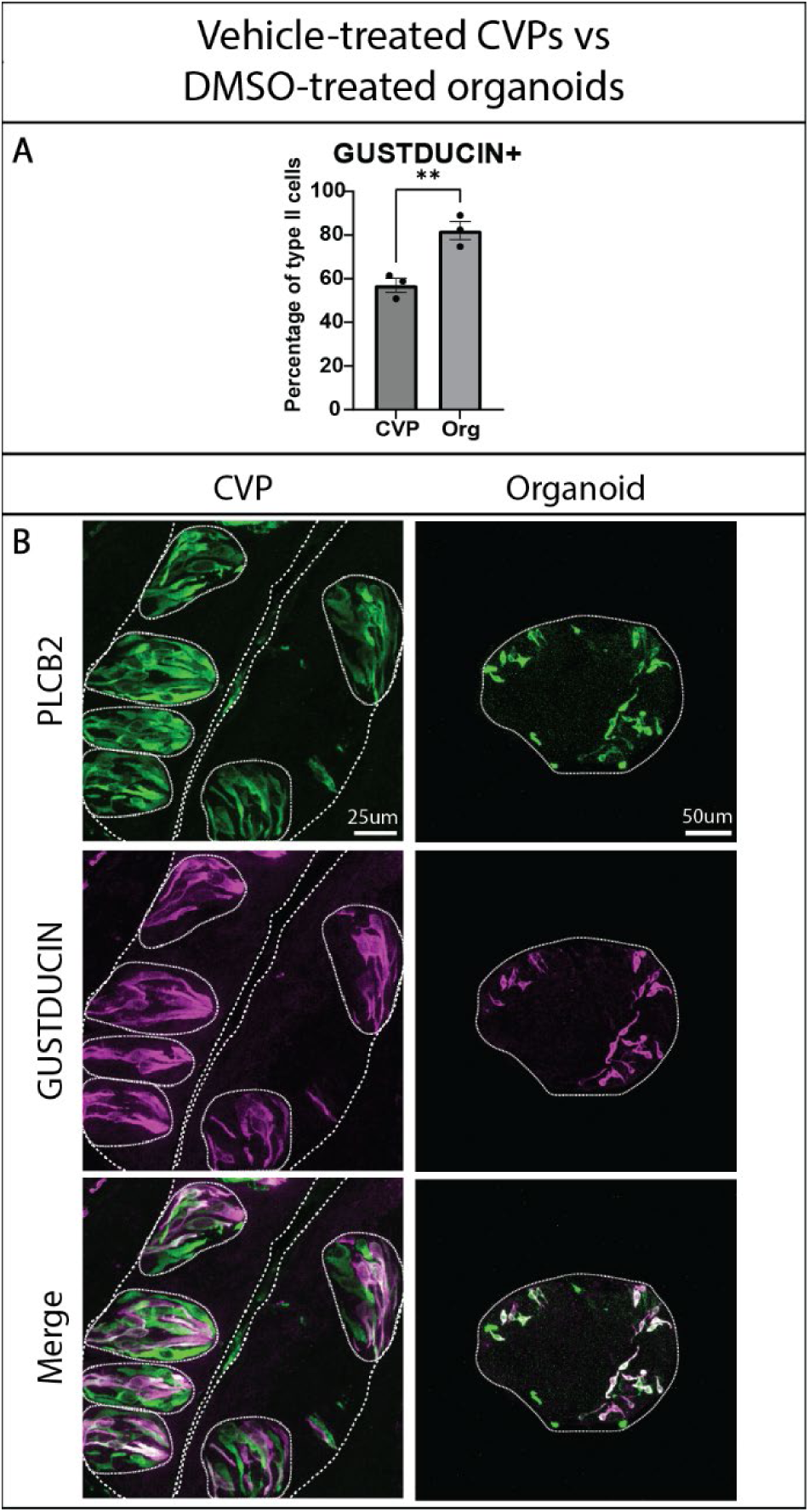
GUSTDUCIN^+^ type II cells are over-represented in lingual organoids compared to CVP taste buds. **(A)** The percentage of PLCβ2^+^ type II cells expressing GUST is higher in organoids compared to *in vivo*. CVP values are from vehicle-treated mice after 4 weeks of treatment (see Figure 2), while organoid values are from organoids treated with DMSO during the differentiation phase (see **Figure 1**). Each dot in the organoid condition is the average percentage across ∼10 organoids for each biological replicate. (**B**) Compressed confocal z-stacks showing vehicle-treated CVP trenches compared to DMSO-treated organoids immunostained for PLCβ2 (green) and GUST (magenta). In the CVP, coarse dashed lines delineate basement membrane and apical surface of epithelium, fine dashed lines encircle individual taste buds. Unpaired t-test. Mean +/- SEM (** p≤0.01).

**Figure S6:**
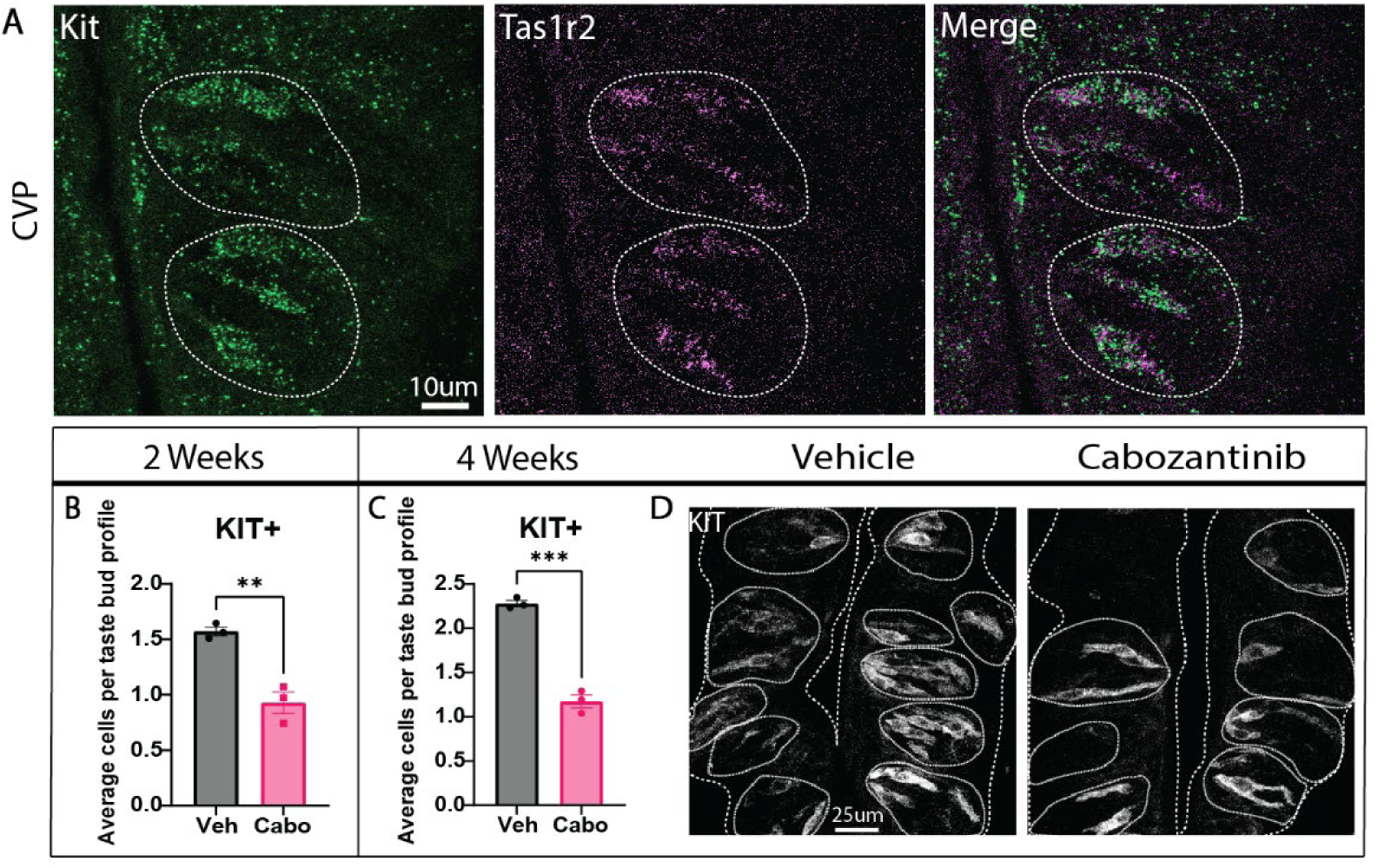
*Kit* marks *Tas1r2*^+^ sweet cells in the CVP. (**A**) In the CVP, HCR *in-situ* hybridization using probes against *Kit* and *Tas1r2* reveals extensive co-expression in taste cells. Dashed lines circle individual taste buds. KIT-IF taste cells per taste bud profile are reduced in the CVP of mice treated with cabozantinib compared to controls at 2 weeks (**B**) and 4 weeks (**C**). (**D**) Compressed confocal z-stacks of KIT immunostaining in mice treated with vehicle or cabozantinib after 4 weeks. Coarse dashed lines delineate basement membrane and apical surface of epithelium, fine dashed lines encircle individual taste buds (N=3 mice per condition). Unpaired t-test. Mean+/-SEM (** p≤0.01, **** p≤0.0001).

**Figure S7:**
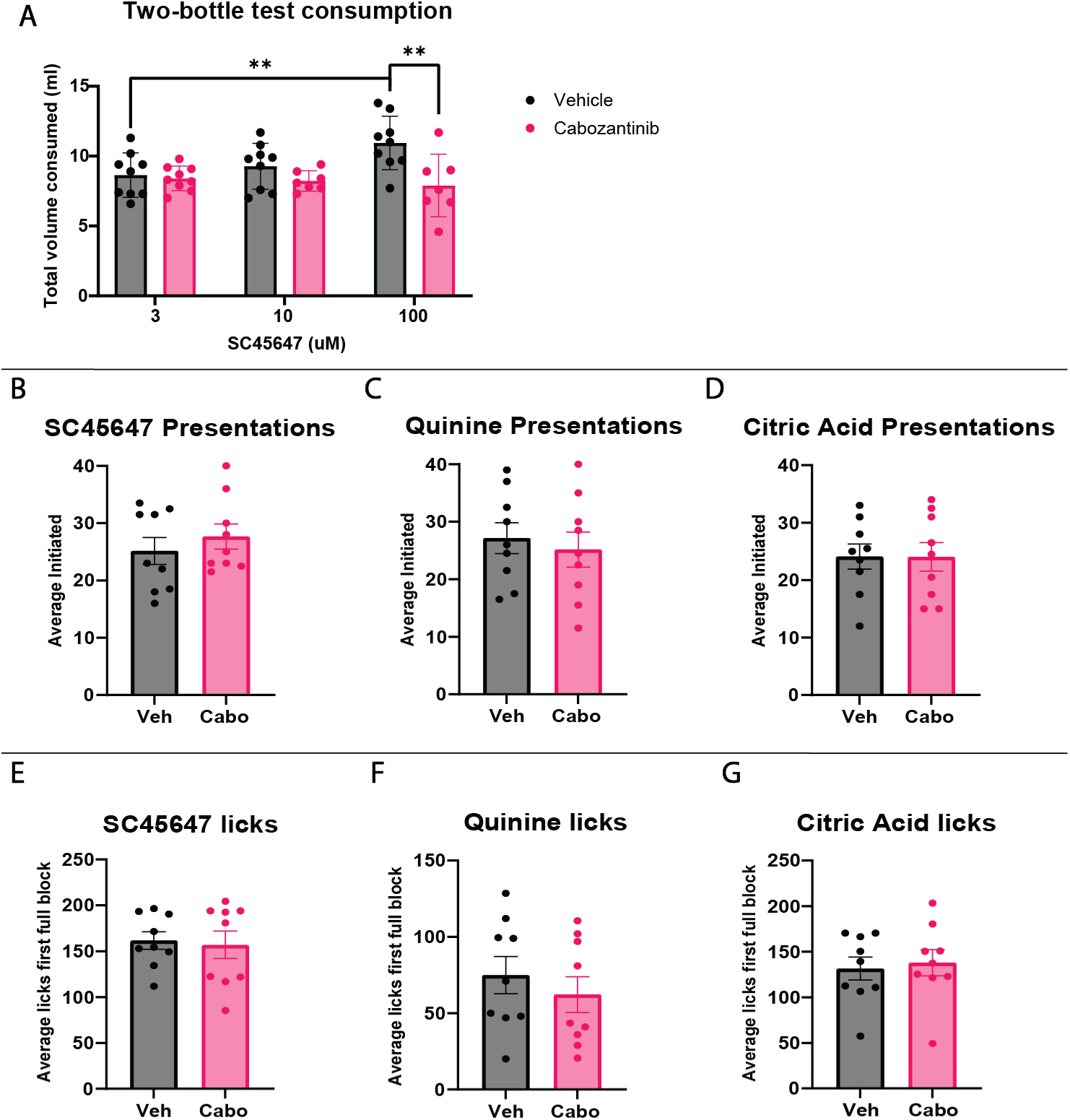
Cabozantinib does not change participation effort or thirst of mice in taste behavior assays. (**A**) Total consumption of liquid across 48 hrs of testing at each tastant concentration did not differ between control and TKI-treated mice, except at the highest concentration of SC45647 where control mice drank significantly more than drug treated mice. (**B-D**) Average presentations initiated across 2 days of lickometer testing for SC45647 (**B**), quinine (**C**), and citric acid (**D**) did not differ with treatment. The maximum number of presentations per testing session was 40. (**E-G**) Average number of licks during the first full testing block (one presentation of each concentration) across 2 testing days did not differ for SC45647 (**E**), quinine (**F**), and citric acid (**G**). Two-way ANOVA with both Sidak’s and Tukey’s multiple comparisons tests performed on data in panel A (N = control 9 and 7 TKI mice). Unpaired t-tests performed on panels B-G (N=9 mice per condition). Mean+/-SEM (** p≤0.01).

**Figure S8:**
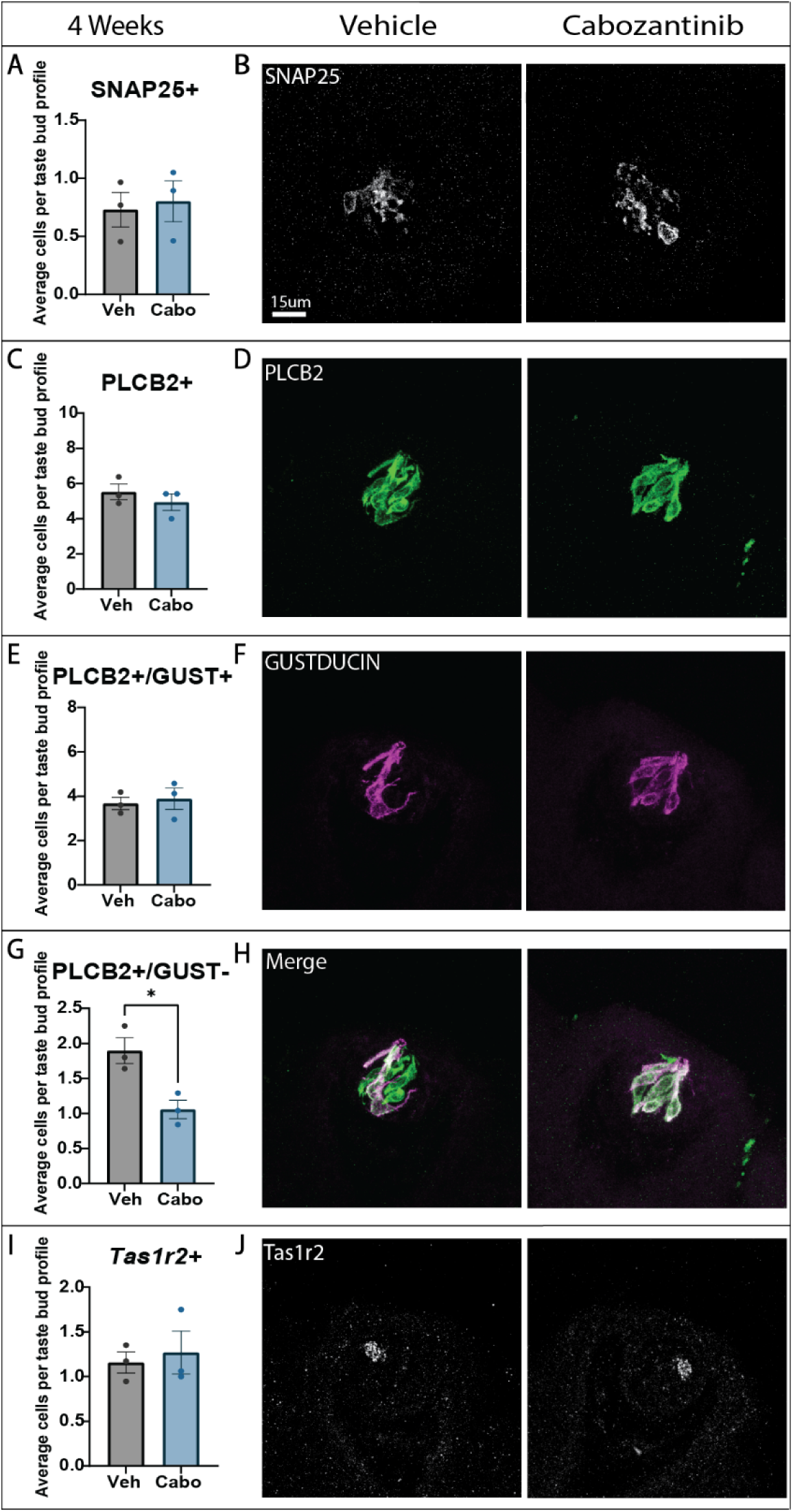
Cabozantinib changes the composition of PLCβ2+ type II subtypes in FFP taste buds. Quantification of the average number of taste cells per taste bud profile with corresponding representative images for SNAP25-IF (**A-B**), PLCβ2-IF (**C-D**), PLCβ2^+^/GUST^+^ IF (**E-F**), PLCβ2^+^/GUST^-^ IF (**G-H**) and *Tas1r2* HCR *in situ* hybridization (**I-J**). Representative images are compressed confocal z-stack projections. Scale bar in B applies to D, F, H and J. In all histograms, each dot represents the average taste cell tally from one mouse (N=3 per condition, ∼10-30 taste buds/mouse). Unpaired t-test. Mean+/-SEM (* p≤0.05).

**Figure S9:**
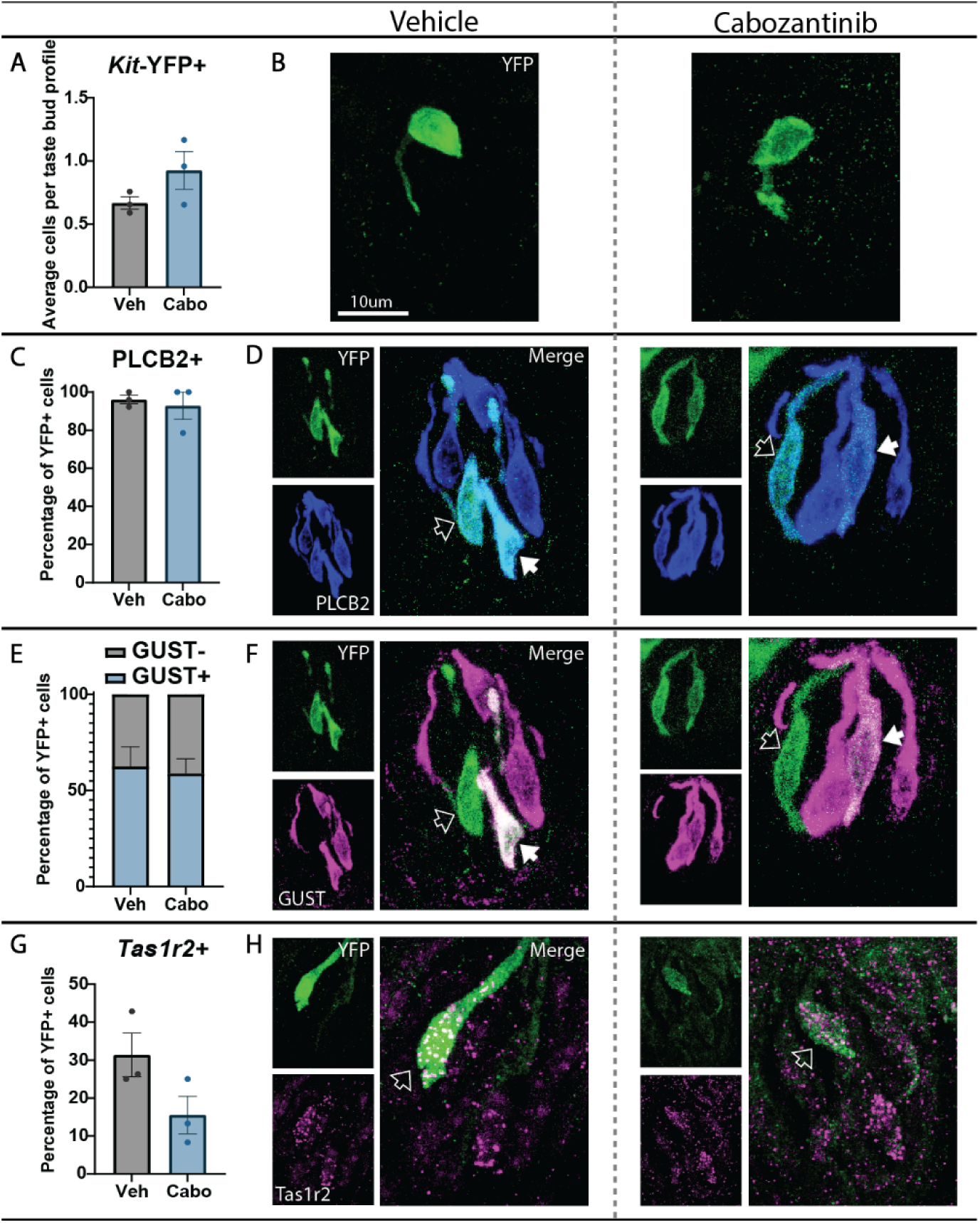
Cabozantinib does not induce cell death or transdifferentiation of *Kit*^+^ cells in FFP taste buds. (**A**) In *Kit^CreER/+^;Rosa26^YFP/YFP^* mice, the average number of *Kit*-YFP^+^ cells per taste bud profile is unaltered by TKI treatment. (**B**) Compressed confocal z-stacks of *Kit*-YFP^+^ taste cells in FFP taste buds from vehicle- vs cabozantinib-treated mice. (**C, E, G**) The percentage of *Kit*-YFP^+^ cells (green in all panels) co-expressing PLCβ2 (**C-D, blue in D**), GUST (**E-F, Magenta in F**) or *Tas1r2* (**G-H, magenta in H**) is unchanged by drug treatment (compressed confocal z-stacks in all panels). Empty arrowheads in **D**, **F** and **H** indicate double-labeled cells, white arrowheads in **D** and **F** indicate triple-labeled cells. In all panels, values were calculated across >50 YFP^+^ cells and >30 taste buds per condition. Unpaired t-test performed for all quantifications. Mean+/-SEM.

**Figure S10:**
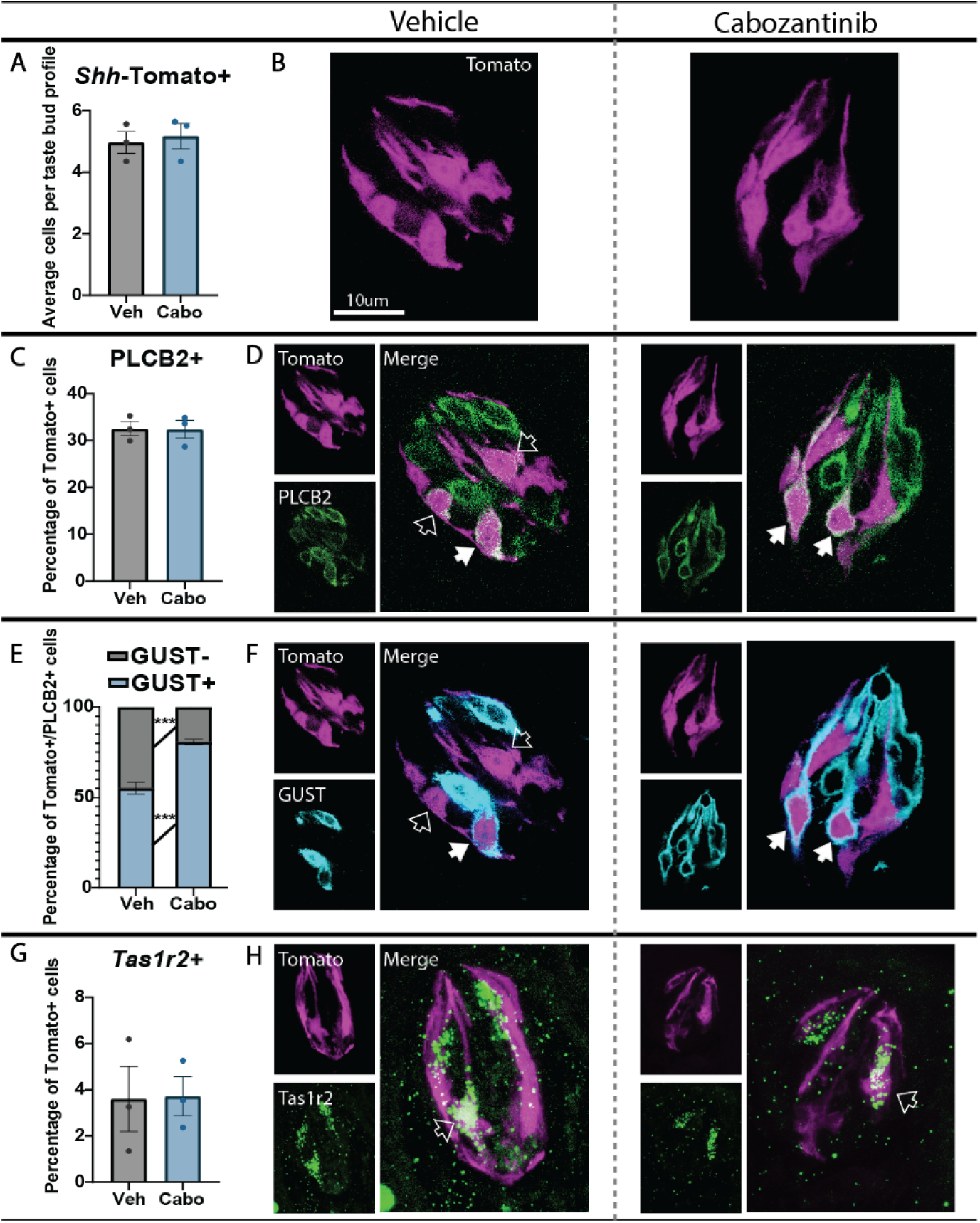
Cabozantinib increases differentiation of PLCß2^+^/GUST^+^ cells in FFP taste buds. (**A**) In *Shh^CreER/+^;Rosa26^tdTomato/+^* mice, the average number of *Shh*-Tomato^+^ cells per taste bud profile was unaltered by cabozantinib treatment. (**B**) Optical section of *Shh*-tomato^+^ taste cells in FFP taste buds from vehicle- vs cabozantinib-treated mice. (**C, E, G**) The percentage of *Shh*-Tomato^+^ cells (**magenta** in all panels) co-expressing PLCβ2 did not change (**C-D, green in D,** optical section). The percentage of *Shh*-Tomato^+^ cells expressing GUST significantly increased and the percentage not expressing GUST significantly decreased (**E-F, cyan in F,** optical section), while the percentage of *Shh*-Tomato^+^ expressing *Tas1r2* did not change (**H-I, green in I,** compressed z-stacks). Empty arrowheads in **D**, **F** and **H** indicate double-labeled cells, white arrowheads in **D** and **F** indicate triple-labeled cells. In all panels, values were calculated across >550 Tomato^+^ cells and >95 taste buds per condition. Unpaired t-test performed for panels **A**, **C** and **G**. Two-way ANOVA with Sidak’s multiple comparisons test performed for panel **E**. Mean+/-SEM (*** p≤0.001).

**Figure S11:**
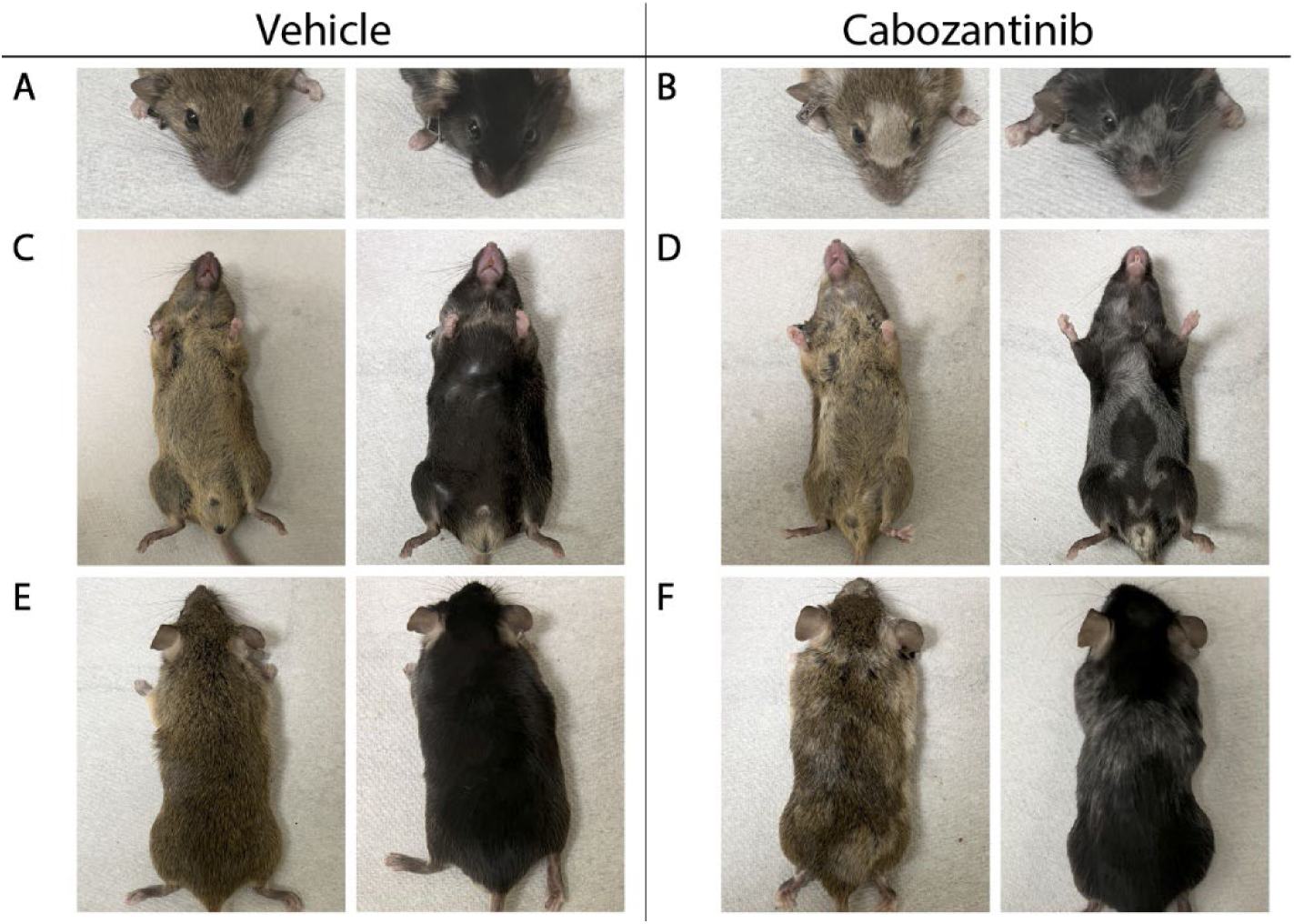
Cabozantinib treatment causes fur depigmentation. Cabozantinib causes depigmentation of facial (**A-B**), stomach (**C-D**) and back (**E-F**) fur. Depigmentation was evident in mice of mixed background with brown or black fur.

**Figure S12:**
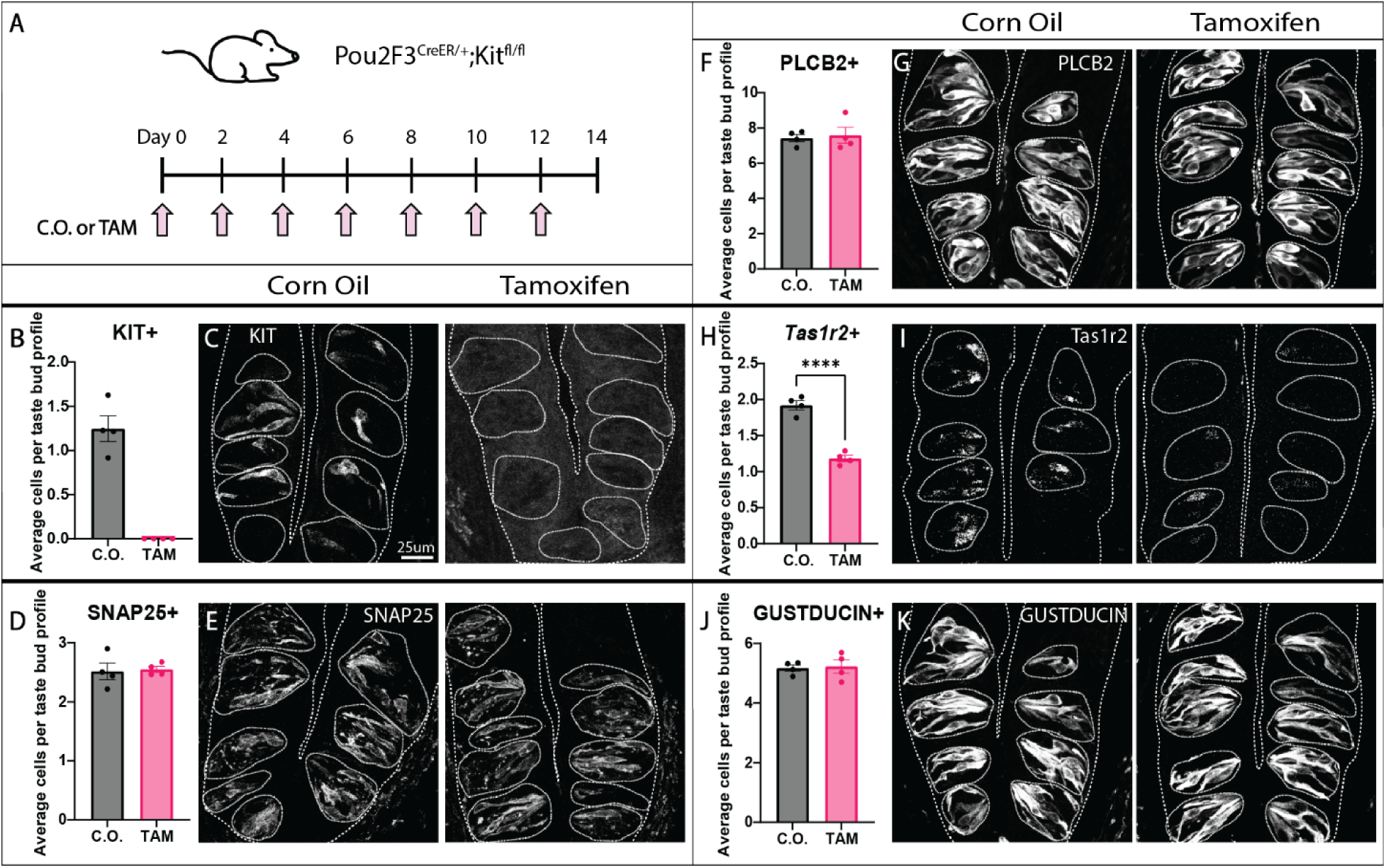
Two-week Kit knockout reduces sweet cells but does not affect bitter/umami cells in CVP. (**A**) *Pou2F3^CreER/+^;Kit ^fl/fl^* mice were gavaged with corn oil or tamoxifen every other day for 14 days. (**B-K**) Quantification of stained cells per taste bud profile with corresponding representative images for KIT-IF (**B-C**), SNAP25-IF (**D-E**), PLCβ2-IF (**F-G)**, *Tas1r2* HCR *in situ* hybridization (**H-I**) and GUST-IF (**J-K**). Representative images are compressed confocal z-stacks. Coarse dashed lines delineate basement membrane and apical epithelial surface, fine dashed lines encircle individual taste buds. Scale bar in **C** applies to **E**, **G**, **I** and **K**. In all histograms, each dot represents the average TBC tally from each mouse (N=4 per condition, ∼80 taste buds/mouse). Unpaired t-test performed on all quantifications. Mean+/-SEM (**** p≤0.0001).

**Figure S13:**
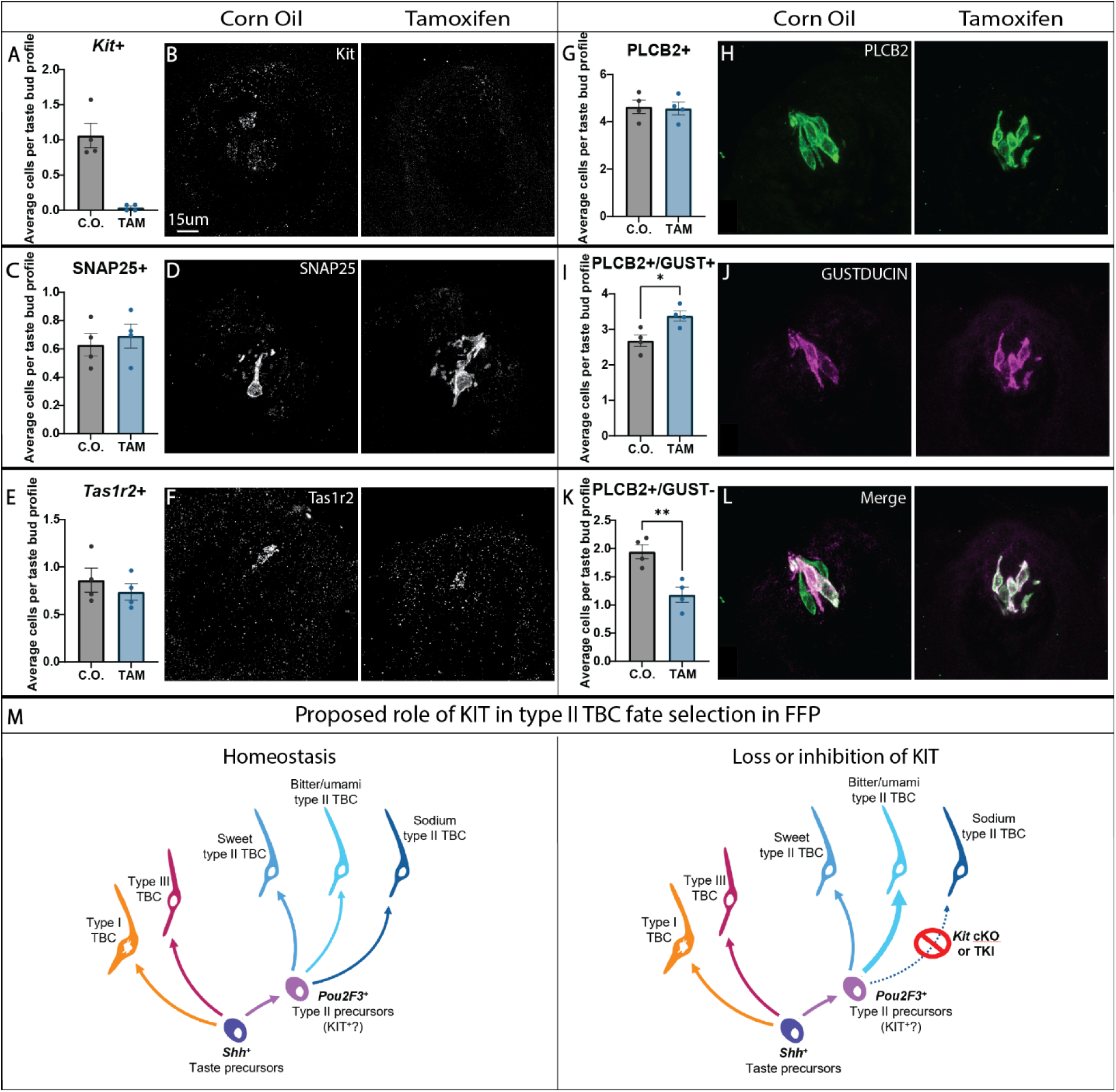
*Kit* knockout causes a fate switch in PLCβ2+ cells in FFP taste buds. (**A-L**) Quantification of the average number of stained cells per taste bud profile with corresponding representative images for *Kit* HCR *in situ* hybridization (**A-B**), SNAP25 IF (**C-D**), *Tas1r2* HCR *in situ* hybridization (**E-F**), PLCβ2 IF (**G-H)**, PLCβ2^+^/GUST^+^ IF (**I-J**), and PLCβ2^+^/GUST^-^ IF (**K-L**). Representative images are compressed confocal z-stack projections. Scale bar in B applies to D, F, H, J and L. In all histograms, each dot represents the average taste cell tally from one mouse (N=4 per condition, ∼15 taste buds/mouse). Mann-Whitney test performed on data from panel A, unpaired t-test performed on data from panels C, E, G, I, K and M. Mean +/- SEM (* p≤0.05, ** p≤0.01). (**M**) Proposed model of *Kit* function in type II TBC lineage. We propose that a *Pou2f3^+^* precursor population gives rise to all type II TBC subtypes. Our pharmacological and genetic data support a model where KIT function is required for sodium-sensing type II cell fate in FFP.

**Table S1:**
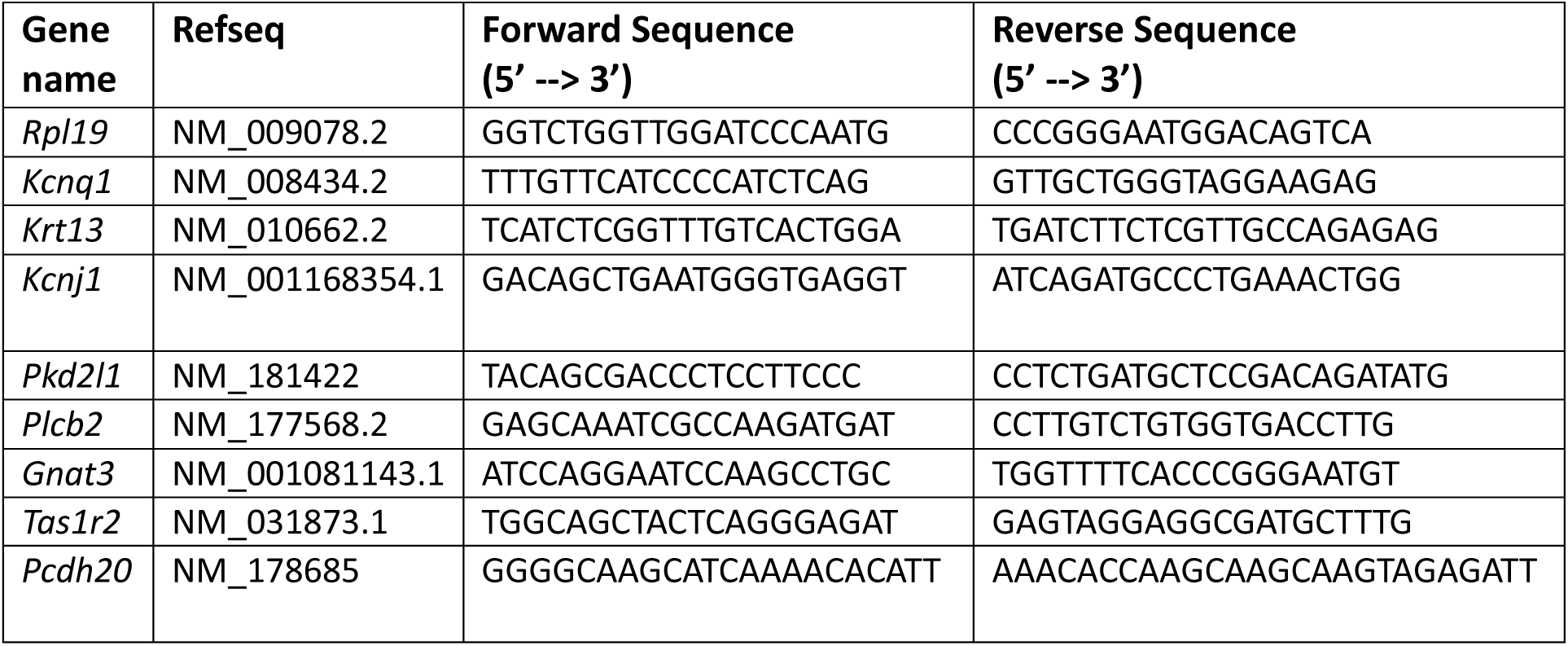
RT-qPCR Primers.

**Table S2:**
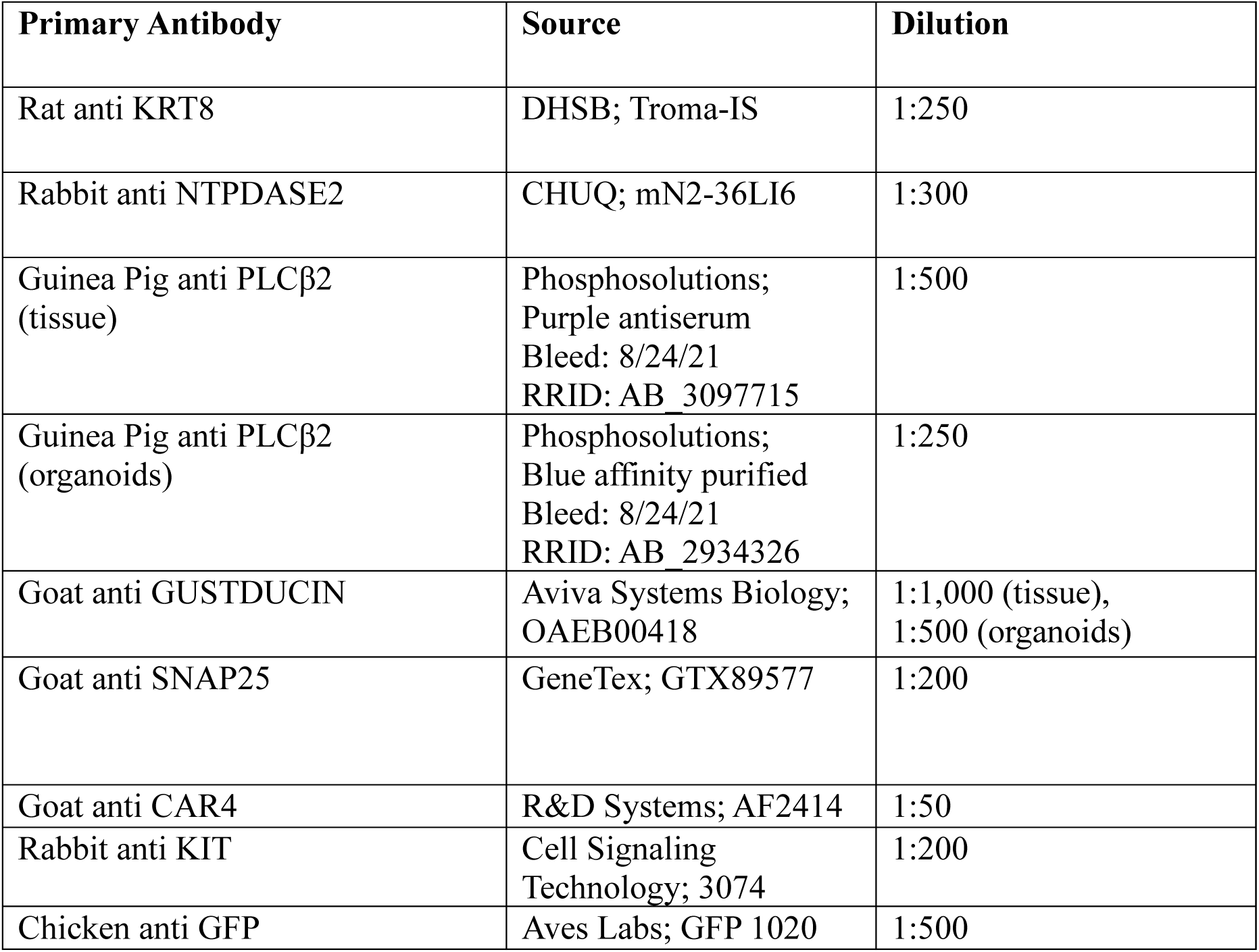
Primary Antibodies.

| **Primary Antibody** | **Source** | **Dilution** |
| --- | --- | --- |
| Rat anti KRT8 | DHSB; Troma-IS | 1:250 |
| Rabbit anti NTPDASE2 | CHUQ; mN2-36LI6 | 1:300 |
| Guinea Pig anti PLCβ2<br>(tissue) | Phosphosolutions;<br>Purple antiserum<br>Bleed: 8/24/21<br>RRID: AB_3097715 | 1:500 |
| Guinea Pig anti PLCβ2<br>(organoids) | Phosphosolutions;<br>Blue affinity purified<br>Bleed: 8/24/21<br>RRID: AB_2934326 | 1:250 |
| Goat anti GUSTDUCIN | Aviva Systems Biology;<br>OAEB00418 | 1:1,000 (tissue),<br>1:500 (organoids) |
| Goat anti SNAP25 | GeneTex; GTX89577 | 1:200 |
| Goat anti CAR4 | R&D Systems; AF2414 | 1:50 |
| Rabbit anti KIT | Cell Signaling<br>Technology; 3074 | 1:200 |
| Chicken anti GFP | Aves Labs; GFP 1020 | 1:500 |

